# Genetic synergy in *Acinetobacter baumannii* undecaprenyl biosynthesis and maintenance of lipid asymmetry impacts outer membrane and antimicrobial resistance

**DOI:** 10.1101/2023.09.22.556980

**Authors:** Hannah R. Noel, Sowmya Keerthi, Xiaomei Ren, Jonathan D. Winkelman, Jerry M. Troutman, Lauren D. Palmer

**Affiliations:** Department of Microbiology and Immunology, University of Illinois Chicago, Chicago, IL, USA; Department of Chemistry, University of North Carolina Charlotte, Charlotte, NC, USA; Trestle LLC, Milwaukee, WI, USA

**Keywords:** *Acinetobacter*, antibiotic resistance, membrane stress, isoprenoid, Und-P, Mla, lipooligosaccharide, LOS

## Abstract

*Acinetobacter baumannii* is a Gram-negative healthcare-associated pathogen that poses a major health concern due to increasing multidrug resistance. The Gram-negative cell envelope is a key barrier to antimicrobial entry and includes an inner and outer membrane. The outer membrane has an asymmetric composition that is important for structural integrity and barrier to the environment. Therefore, Gram-negative bacteria have mechanisms to uphold this asymmetry such as the maintenance of lipid asymmetry system (Mla), which removes glycerophospholipids from the outer leaflet of the outer membrane and transports them to the inner membrane. Loss of this system in *A. baumannii* results in attenuated virulence and increased susceptibility to membrane stressors and some antibiotics. We recently reported two strain variants of the *A. baumannii* type strain ATCC 17978, 17978VU and 17978UN. We show here that Δ*mlaF* mutants in the two strains display different phenotypes for membrane stress resistance, antibiotic resistance, and pathogenicity in a murine pneumonia model. We used comparative genetics to identify interactions between ATCC 17978 strain alleles and *mlaF* to uncover the cause behind the phenotypic differences. Although allele differences in *obgE* were previously reported to synergize with Δ*mlaF* to affect growth and stringent response, we show that *obgE* alleles do not affect membrane stress resistance. Instead, a single nucleotide polymorphism (SNP) in the essential gene encoding undecaprenyl pyrophosphate (Und-PP) synthase, *uppS*, synergizes with Δ*mlaF* to increase susceptibility to membrane stress and antibiotics, and reduce persistence in a mouse lung infection. Und-P is a lipid glycan carrier known to be required for biosynthesis of *A. baumannii* capsule, cell wall, and glycoproteins. Our data suggest that in the absence of the Mla system, the cellular level of Und-P is critical for envelope integrity, antibiotic resistance, and lipooligosaccharide abundance. These findings uncover synergy between Und-P and the Mla system in maintaining the *A. baumannii* outer membrane and stress resistance.

## Introduction

*Acinetobacter baumannii* is a Gram-negative bacterial pathogen that is a major cause of hospital-acquired infection. Many clinical isolates of *A. baumannii* demonstrate resistance to first line antibiotics such as meropenem and colistin (1, 2). Both The World Health Organization and the Centers for Disease Control have thus identified *A. baumannii* as an urgent threat, calling for the development of novel antimicrobials (3–5). *A. baumannii* has multiple intrinsic mechanisms for resisting antibiotics and host stressors. The first line of defense is the cellular envelope including capsule, a peptidoglycan cell wall, and a dual-membrane system conserved among Gram-negative bacteria (6, 7). The inner membrane (IM) and outer membrane (OM) membrane protect the bacteria from environmental stress (8, 9). The Gram-negative OM is asymmetric with phospholipids composing the inner leaflet and lipopolysaccharides (LPS) or lipooligosaccharides (LOS) composing the outer leaflet. *A. baumannii* does not encode the gene required for O-antigen elaboration as other Gram-negative bacteria do, resulting in LOS rather than LPS (10). Additionally, *A. baumannii* is able to survive without LOS, whereas in most Gram-negative bacteria, LPS is essential (11, 12). In all Gram-negative bacteria, the asymmetric bilayer of the OM is critical for resistance to membrane stressors and antimicrobials (13).

The maintenance of lipid asymmetry (Mla) system is thought to be the primary homeostatic mechanism to maintain OM lipid asymmetry by removing mislocalized phospholipids from the outer leaflet of the OM (14, 15). MlaBDEF forms an IM ATP binding cassette (ABC) transporter with ATPase activity, MlaC is a periplasmic protein, and MlaA is an OM lipoprotein that forms a complex with OmpC/F (14, 16–18). Disruption of the Mla system results in increased outer membrane permeability and drug sensitivity (14, 19). Additionally, dominant negative mutations in *mlaA* (*mlaA*\*) have been shown to increase outer membrane permeability and sensitivity to erythromycin and rifampicin in *Escherichia coli* (19). While the Mla system is not required for growth in lysogeny broth, bacteria lacking a functional Mla system are more sensitive to the membrane stress SDS/EDTA (14, 16, 20–22). Inactivating *mlaF* or *mlaC* in multiple species results in the loss of function of the Mla system and increased sensitivity to membrane stressors, antibiotics, and the host (20, 23–25). The Mla system is critical for the virulence of multiple pathogenic species including *Shigella flexneri*, *Burkholderia pseudomallei*, and *Pseudomonas aeruginosa* (26–30). By contrast, mutations in the Mla system have been shown to increase virulence in *E. coli* and *Neisseria gonorrhoeae* (31, 32). In summary, the Mla system is critical for maintenance of the gram-negative outer membrane, stress resistance, and virulence.

While the directionality of lipid transport by the Mla system has been debated, the preponderance of evidence support a retrograde transport model of phospholipid movement from the OM to the IM (33–35). Genetic evidence from *E. coli, A. baumannii,* and chloroplasts support a retrograde transport model in which phospholipids are removed from the outer leaflet of the OM and transported to the IM (14, 19, 20, 36–39). Additionally, crystallographic and cryo-EM structural data from *Klebsiella pneumonia*, *Serratia marcescens,* and *E. coli* support a model in which phospholipids are removed from the OM by the MlaA-OmpF complex and transported towards MlaBDEF based on the orientation of MlaA-OmpF in the OM (16, 40–42). Recent studies identifying AsmA-like proteins facilitating anterograde lipid transport machinery in *E. coli* further support the retrograde transport model for the Mla system (43, 44). By contrast, *in vitro* work in *E. coli* as well as cryo-EM, molecular dynamics and pulse-chase studies in *A. baumannii* described an anterograde transport model where newly synthesized phospholipids are transported to the inner leaflet of the OM (18, 23, 45, 46). However, Mann *et. al.*, whose structural work in *A. baumannii* supports a lipid export model, speculate that nucleotide state or solubilization approach of the MlaBDEF studies may explain the differences in conclusions (46). Recently, structures of MlaBDEF from *E. coli* and *A. baumannii*, and *E. coli* MlaC in complex with MlaA or MlaD were solved and provide mechanistic insight on lipid binding, but do not provide clear evidence for either direction of lipid transport (21, 47, 48).

In *A. baumannii*, the Mla system synergizes with essential pathways to promote growth. A previous study reported synergy between the Mla system and the essential GTPase ObgE in promoting Δ*mlaF* growth and stringent response (49, 50). We previously reported a suppressor mutation in the isoprenoid biosynthetic pathway that restored resistance of an *A. baumannii* Δ*mlaF* mutant to membrane stress, some antibiotics, and host stressors (20). The suppressor is an IS*Aba11* transposition in the 5’ untranslated region of *ispB* that results in *ispB* downregulation (20). We hypothesized that the downregulation of *ispB* increased flux of the branchpoint substrate farnesyl-pyrophosphate (FPP) to undecaprenyl pyrophosphate (Und-PP) synthase, UppS. Undecaprenyl pyrophosphate synthase produces Und-PP, which is a precursor to the essential glycan carrier undecaprenyl phosphate (Und-P) (51). Und-P is responsible for transferring glycans across the plasma membrane to produce important envelope components like peptidoglycan, capsular polysaccharides, and the O-antigen in LPS-producing bacteria. This finding suggested a role for Und-P biosynthesis in membrane stress resistance in the absence of the Mla system. Thus, multiple reports have identified synergy between the Mla system and other pathways in *A. baumannii*.

Recently, we identified two variants of the commonly used laboratory strain *A. baumannii* 17978 distributed by ATCC (52). These variants, *A. baumannii* ATCC 17978VU and 17978UN, have distinct genotypes with 6 protein-encoding single nucleotide polymorphisms (SNPs) as well as a 44 kb accessory locus (AbaAL44) that is present in 17978UN but absent in 17978VU (52). The protein-coding SNPs encode variants of the predicted essential proteins ObgE, UppS, and the lipooligosaccharide transporter LptD (53). Our previous work used isogenic Δ*mlaF* strains in the 17978UN (AbaAL44^+^) background (20). Upon reconstructing Δ*mlaF* strains in the *A. baumannii* ATCC 17978VU strain background, we discovered that the *ΔmlaF* mutant was more resistant to SDS/EDTA membrane stress in the 17978VU background compared to the 17978UN background. Here we describe genetic synergy between the maintenance of lipid asymmetry and Und-P biosynthesis uncovered through genetic dissection and comparison of ATCC 17978VU and 17978UN Δ*mlaF* strains. These findings suggest an underlying relationship between the Mla system and undecaprenyl biosynthesis in *A. baumannii* that function together to maintain LOS abundance and promote membrane stress resistance, antimicrobial resistance, and virulence.

## Results

### UppS synergizes with the Mla system under membrane stress

The 17978VU and 17978UN strain variants of ATCC 17978 contain SNPs in multiple protein coding genes (52). However, it is unknown which variant contains the more representative alleles for *A. baumannii* strains in general. To address this, we assessed the prevalence of the SNP containing genes across *A. baumannii* published genomes. A set of 5945 genomes were de-duplicated to remove near clonal lineages and the predicted proteomes of the resulting 233 genomes were further analyzed by OrthoFinder to deduce orthologues within *A. baumannii* (54, 55). The 17978UN predicted protein variants were more common for LptD (99%) and ClsC2 (indicative of the presence of the AbaAL44 cluster; 90%). The 17978VU predicted protein variants were more common for ObgE (99%), UppS (99%), ActP (99%), the amino acid symporter (79%), and DUF817 (89%) (Fig. 1). Thus, both 17978VU and 17978UN contain alleles representing the majority of published *A. baumannii* genomes. We reconstructed the Δ*mlaF*::Kn (Δ*mlaF*) mutation in *A. baumannii* ATCC 17978VU and compared SDS/EDTA membrane stress resistance in Δ*mlaF* in the 17978VU (AbaAL44^-^) and 17978UN (AbaAL44^+^) backgrounds by growth curves in LB with and without SDS/EDTA. Neither the wild-type strains nor the Δ*mlaF* mutants demonstrated a growth defect in LB alone regardless of strain background (Fig. 2A and 2B). However, in the presence of membrane stress, the 17978UN Δ*mlaF* mutant exhibited a greater growth defect than the 17978VU Δ*mlaF* mutant (Fig. 2F and 2G), suggesting synergy between *mlaF* and genetic differences between the strains. Therefore, the closely related strain variants ATCC 17978VU and 17978UN could be used as a tool to uncover this synergy.

**Figure 1.**
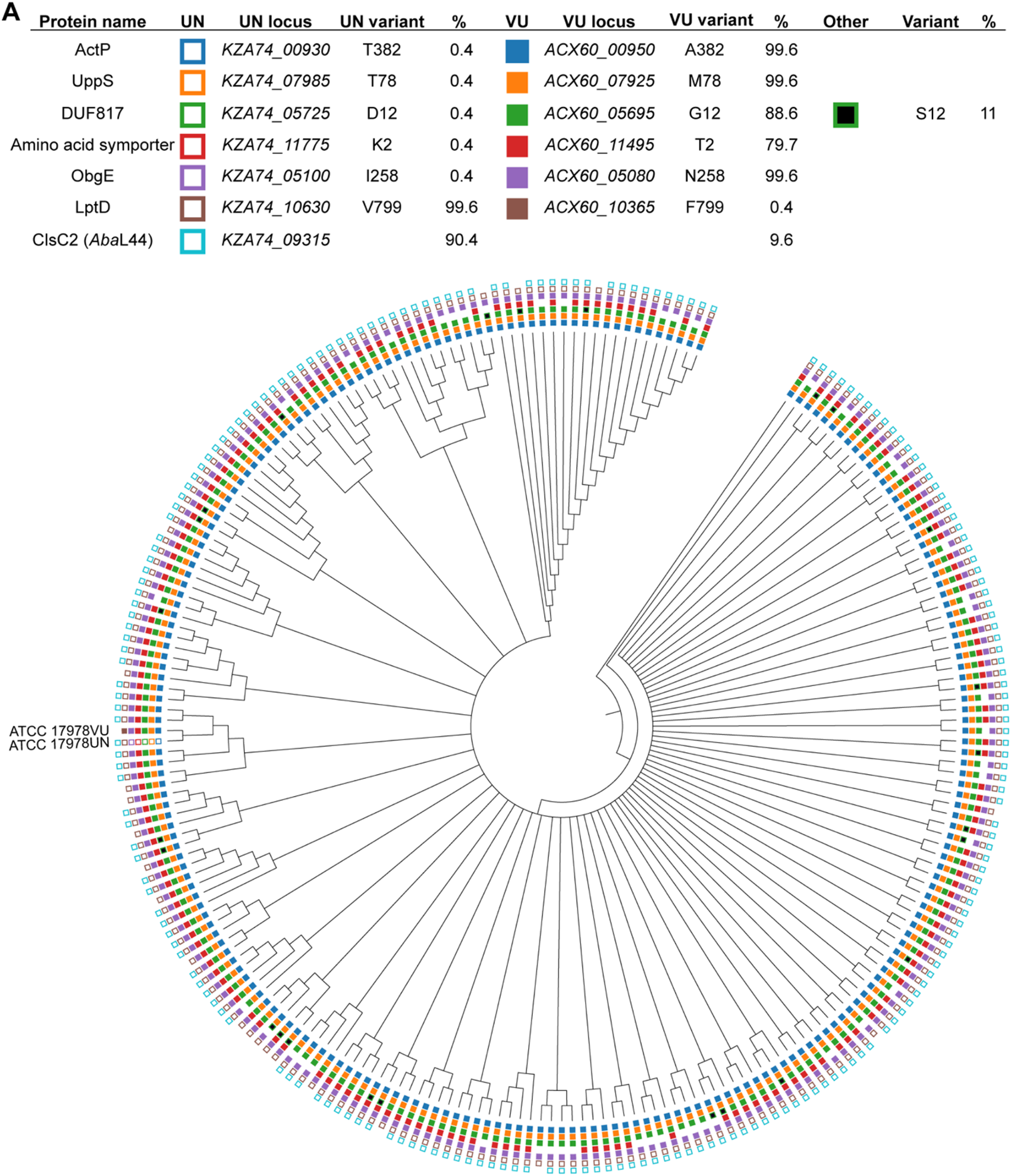
Phylogenetic tree of *Acinetobacter baumannii* strains depicting prevalence of variants encoded by 17978VU and 17978UN. Filled, non-black boxes indicate the presence of a protein variant identical to *A. baumannii* 17978VU, while unfilled boxes denote the presence of the *A. baumannii* 17978UN variant. The absence of a box indicates that an ortholog was not found in the genome. Black boxes indicate the presence of a variant different from both the VU and UN strains. The presence of ClsC2, indicated by an unfilled box of the 17978UN strain, is representative of the presence of the 44-kb accessory locus AbaAL44.

**Figure 2.**
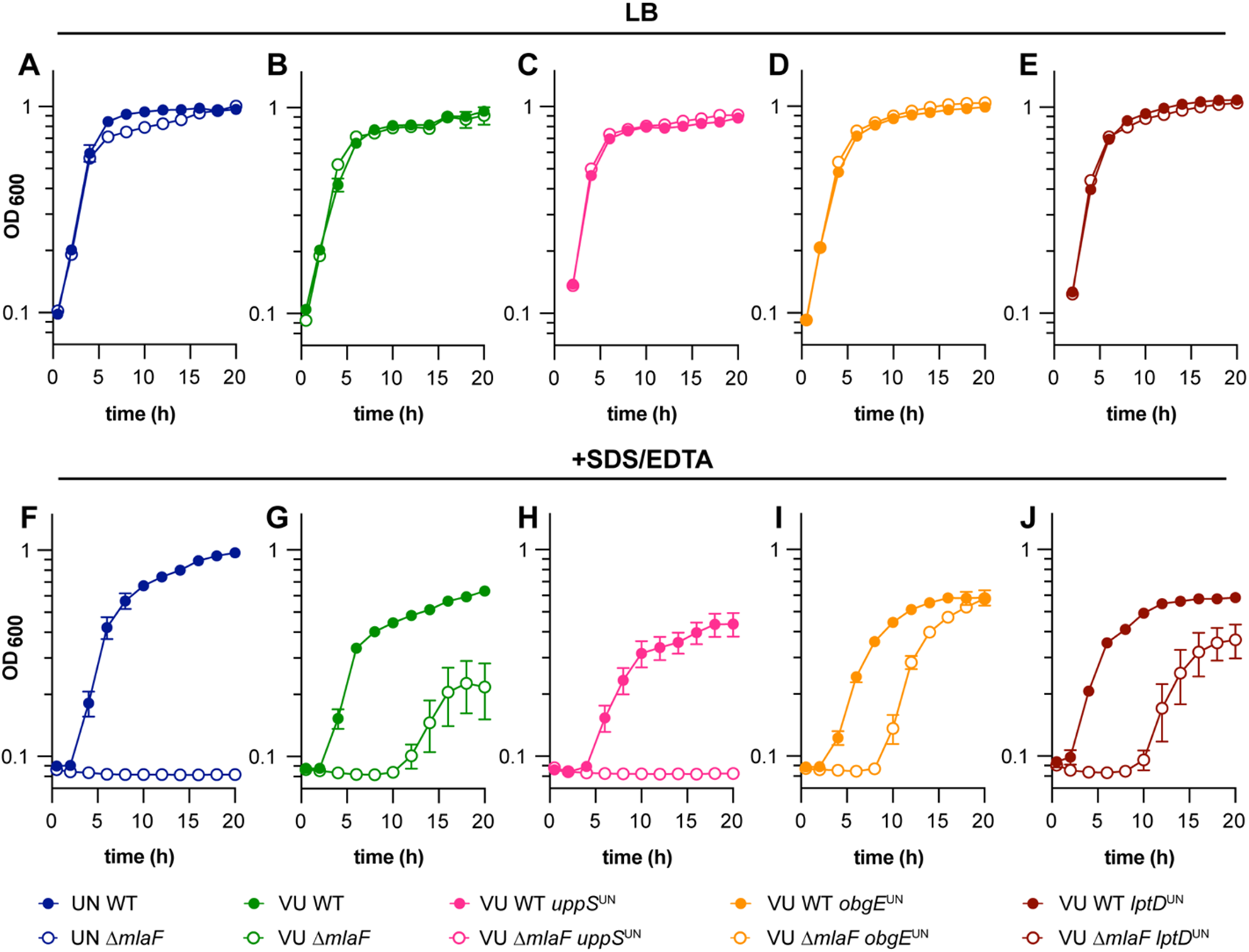
UppS^UN^ protein variant results in increased membrane stress sensitivity in *A. baumannii* ATCC 17978 Δ*mlaF* mutants. **(A to E)** ATCC 17978 UN and VU WT, Δ*mlaF,* and isogenic mutant strains were grown in LB. **(F to J)** ATCC 17978 UN and VU WT, Δ*mlaF,* and isogenic mutant strains were grown in LB with 0.01% SDS and 0.175 mM EDTA. Data are means ± SEM, n=3.

To determine the genetic cause behind the differences in membrane stress resistance between ATCC 17978VU and 17978UN Δ*mlaF* strains, point mutants were constructed for three candidate genes: *obgE*, *lptD*, and *uppS*. Candidate genes were chosen based on literature findings and known function. First, Powers *et al*. previously reported that *obgE* alleles in 17978 strains maintained at University of Washington (UW) and University of Georgia (UGA) synergized with *mlaF* to affect growth and stringent response. Specifically, the Δ*mlaF* strain with ObgE^I258^ (ObgE^UN^) had defects in LB growth and stringent response compared to the Δ*mlaF* strain with ObgE^N258^ (ObgE^VU^)(49). The strains were otherwise isogenic, suggesting that one strain was a derivative of 17978VU or 17978UN. Second, LptD is a β-barrel OM protein responsible for the translocation of LPS/LOS in Gram-negative bacteria (56). Third, we previously reported a potential role for undecaprenyl pyrophosphate, synthesized by *uppS*, in promoting *A. baumannii* envelope integrity in 17978UN Δ*mlaF* (20). The 17978UN allele for each candidate gene (encoding ObgE I258; LptD V799; UppS T78) was substituted in the intrinsic locus (encoding ObgE N258; LptD F799; UppS M78) in 17978VU WT and 17978VU Δ*mlaF* strains. We reasoned that if the candidate gene contributes to the contrasting phenotypes, then the 17978VU Δ*mlaF* mutant containing the 17978UN candidate allele would exhibit the 17978UN Δ*mlaF* phenotype of increased membrane stress sensitivity. As expected, none of the strains display a growth defect when grown in LB alone (Fig. 2A-E). However, in the presence of SDS/EDTA, neither the *obgE* nor *lptD* 17978UN alleles increased membrane stress sensitivity in the 17978VU Δ*mlaF* strain. Rather, 17978VU Δ*mlaF uppS^UN^* was unable to grow in SDS/EDTA similarly to 17978UN Δ*mlaF*, suggesting that the *uppS* allele determines membrane stress sensitivities of 17978VU and 17978UN Δ*mlaF* strains (Fig. 2H). Together, these data show that the Mla system is synthetic with *uppS* alleles in *A. baumannii* for resistance to SDS/EDTA membrane stress.

### uppS alleles influence membrane permeability

Given the role of both UppS and the Mla system in cell envelope biosynthesis and integrity, we hypothesized that other OM-associated phenotypes would be impacted by the *uppS* alleles in a Δ*mlaF* mutant. We predicted that the *uppS*^UN^ allele would be associated with increased membrane permeability. An ethidium bromide (EtBr) uptake assay was used to test the effect of the *uppS* alleles on membrane permeability in wild-type and Δ*mlaF* strains (23, 57). In the 17978VU background, Δ*mlaF uppS*^UN^ has significantly increased EtBr permeability compared to wild type, Δ*mlaF,* and *uppS*^UN^ (Fig. 3A, C). Likewise, in the 17978UN background, introducing the *uppS*^VU^ reduces membrane permeability in an Δ*mlaF* mutant (Fig. 3B, D). Together, these data suggest that *uppS* and the Mla system synergize to promote membrane integrity.

**Figure 3.**
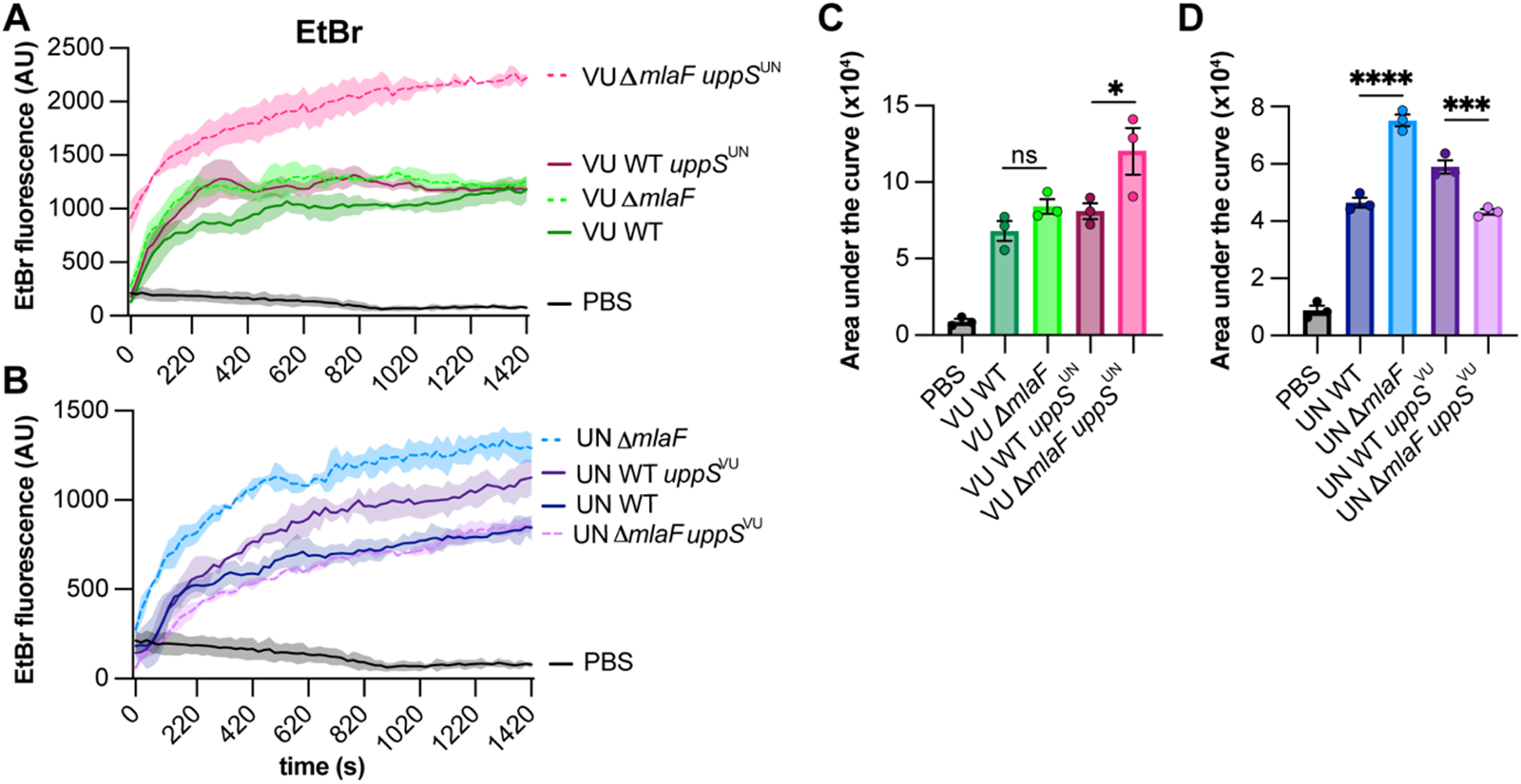
UppS^UN^ increases membrane permeability in 17978 Δ*mlaF* strain. **(A and B)** ATCC 17978VU (A) and 17978UN (B) WT and Δ*mlaF* strains with the intrinsic or opposite *uppS* allele were incubated with efflux pump inhibitor CCCP and ethidium bromide and fluorescence was measured. Data are representative of two experiments. n=3, data are means ± SEM. **(C and D)** Quantified area under the curve for A and B, respectively. Significance is by one-way ANOVA with Tukey’s multiple comparisons test. Comparisons were made each strain and the PBS-only negative control and between Δ*mlaF* strains with their respective wildtype strain. significant differences are indicated by * *p* < 0.05, ** *p* <0.01, *** *p* < 0.001, **** *p* < 0.0001.

### Isoprenoid pathway mutations suppress ΔmlaF membrane stress sensitivity only in the presence of uppS^UN^

To determine if the *uppS*^VU^ allele would result in increased membrane stress resistance in a 17978UN Δ*mlaF* background, the *uppS*^VU^ allele was introduced into 17978UN WT and 17978UN Δ*mlaF* strains. Our previously identified 17978UN Δ*mlaF* suppressor was included to test the contribution of the isoprenoid biosynthesis pathway to the contrasting phenotypes (outlined in Fig. 4A). The suppressor mutation, which decreases transcript abundance of *ispB*, was identified in a 17978UN Δ*mlaF* background and restored virulence and membrane stress and antibiotic resistance to wildtype levels (20). Thus, to determine if the isoprenoid biosynthetic pathway impacts Δ*mlaF* growth, 17978UN Δ*mlaF ispB::*IS*Aba11* and 17978VU Δ*mlaF ispB::*IS*Aba11* strains were constructed with the intrinsic or alternate *uppS* allele. We performed a dilution spotting experiment to assess colony formation efficiency. As expected, none of the strains exhibit defects in colony formation on LB alone (Fig. 4B). In the presence of SDS/EDTA membrane stress, Δ*mlaF* strains encoding UppS^UN^ exhibited a clear defect in colony formation efficiency compared to Δ*mlaF* strains encoding UppS^VU^ (Fig. 4C). This further supports the conclusion of *uppS* being the genetic cause behind the differences in membrane stress resistance between Δ*mlaF* strains in ATCC 17978VU and 17978UN. Similarly, the *ispB* suppressor restored colony formation on SDS/EDTA in Δ*mlaF* strains encoding the *uppS*^UN^ allele, regardless of strain background (Fig. 4C). This suggests that the *ispB* suppressor function is associated specifically with *uppS*^UN^.

**Figure 4.**
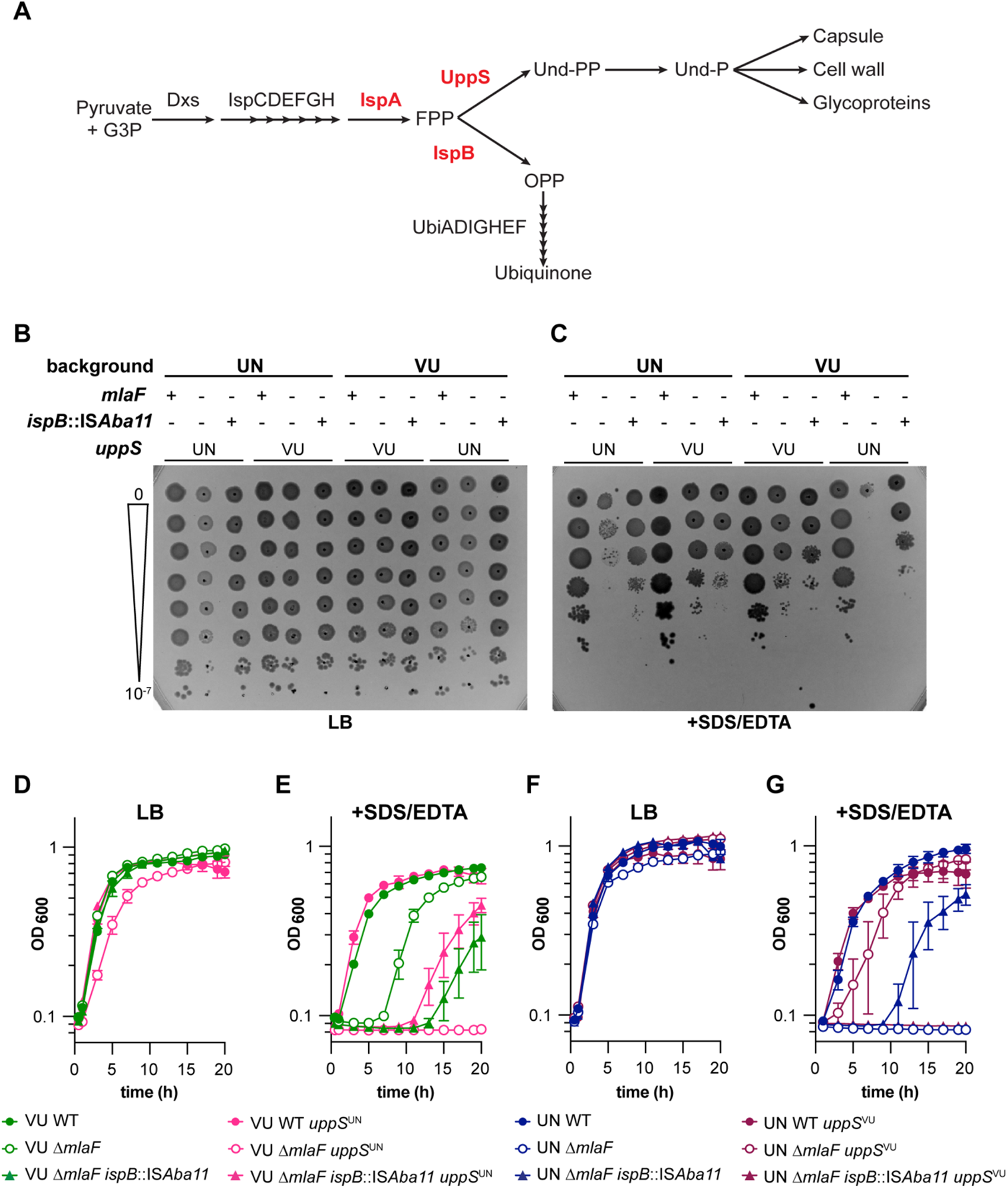
Isoprenoid biosynthesis suppressor mutations require UppS^UN^ to confers increased resistance to SDS/EDTA in 17978 Δ*mlaF* strains. **(A)** Schematic of the isoprenoid biosynthetic pathway with key genes highlighted in red. **(B and C)** WT and mutant strains were serially diluted before spotting on LB plates without (**B**) and with (**C**) 0.01% SDS and 0.15 mM EDTA. Data are representative of 4 experiments with a total of 8 biological replicates. **(D to G)** VU (D and E) and UN (F and G) WT and mutant strains were grown in LB without (D and F) or with (E and G) 0.01% SDS and 0.175 mM EDTA. n=3, data are means ± SEM.

To examine the effect of the different *uppS* alleles on Δ*mlaF* growth over time, we performed growth curves in liquid media with or without SDS/EDTA. Again, none of the strains showed overt growth defects when grown in LB alone (Fig. 4D, 4F). However, in the Δ*mlaF* background, the *uppS*^UN^ allele resulted in complete lack of growth in media containing SDS/EDTA. Conversely, the *uppS*^VU^ allele conferred increased resistance to SDS/EDTA in the Δ*mlaF* background (Fig. 4E, 4G). Interestingly, 17978UN Δ*mlaF ispB::*IS*Aba11 uppS*^VU^ was unable to grow in the presence of membrane stressors SDS/EDTA (Fig. 4G). This result suggests that the *uppS*^VU^ allele in a 17978UN Δ*mlaF* background may be hindered by the *ispB* suppressor, perhaps due to a decreased flux towards ubiquinone production.

While constructing the 17978VU Δ*mlaF uppS*^UN^ strain, one isolate displayed a distinct phenotype resembling that of 17978VU Δ*mlaF*. Upon whole genome sequencing, a mutation was discovered in *ispA* that resulted in a G223E substitution. This mutation was then reconstructed in the 17978UN Δ*mlaF* background to assess its function as a suppressor of the Δ*mlaF uppS*^UN^ phenotype. A dilution spotting assay was used to characterize both *ispA* and *ispB* suppressors in the 17978UN background. All four strains grew well on LB medium (Fig. S1). As expected, 17978UN Δ*mlaF* displayed a strong growth defect that is partially rescued by suppressor mutations in *ispA* and *ispB* (Fig. S1). These data support a model in which the isoprenoid biosynthetic pathway is synthetic with the Mla system in *A. baumannii*. We predict that the mechanism in which both Δ*mlaF* suppressors function is by increasing the metabolic flux towards UppS and the production of Und-P, which promotes growth in the presence of the *uppS*^UN^ allele.

### UppS^UN^ has decreased enzymatic activity, leading to lower cellular Und-P levels and LOS abundance

Thus far, the data suggested that there is a difference in function between UppS^VU^ and UppS^UN^. We hypothesized that UppS^UN^ (T78) was defective compared to UppS^VU^ (M78). First, secondary structure was compared by circular dichroism analysis which showed that there were no structural differences between the UppS variants (Fig. S2A-B). To compare enzymatic activity, enzyme assays were performed using purified protein and a fluorescent analog (2-nitrileanilinogeranyl diphosphate; 2CNA-GPP) of the UppS substrate farnesyl pyrophosphate. UppS from *Escherichia coli, Vibrio vulnificus, Staphylococcus aureus,* and *Bacteroides fragilis* have previously been shown to catalyze the extension of the fluorescent substrate analogue 2CNA-GPP (58, 59). Upon elongation of the analogue, a concomitant increase in fluorescence can be readily monitored via a microplate assay. By this assay, UppS^UN^ was found to have a five-fold decrease in the rate of fluorescence signal compared to UppS^VU^ (Fig 5A). To determine if the decreased enzymatic activity of UppS^UN^ results in decreased levels of the essential glycan carrier Und-P in *A. baumannii*, we quantified the cellular pools of Und-P in the 17978VU and 17978UN wild-type strains. The 17978UN strain contained two-fold lower levels of Und-P compared to the 17978VU wild-type strain (Fig. 5B). These data suggest that the UppS^UN^ enzyme is aberrant in function and has insufficient activity to maintain cellular levels of Und-P.

**Figure 5.**
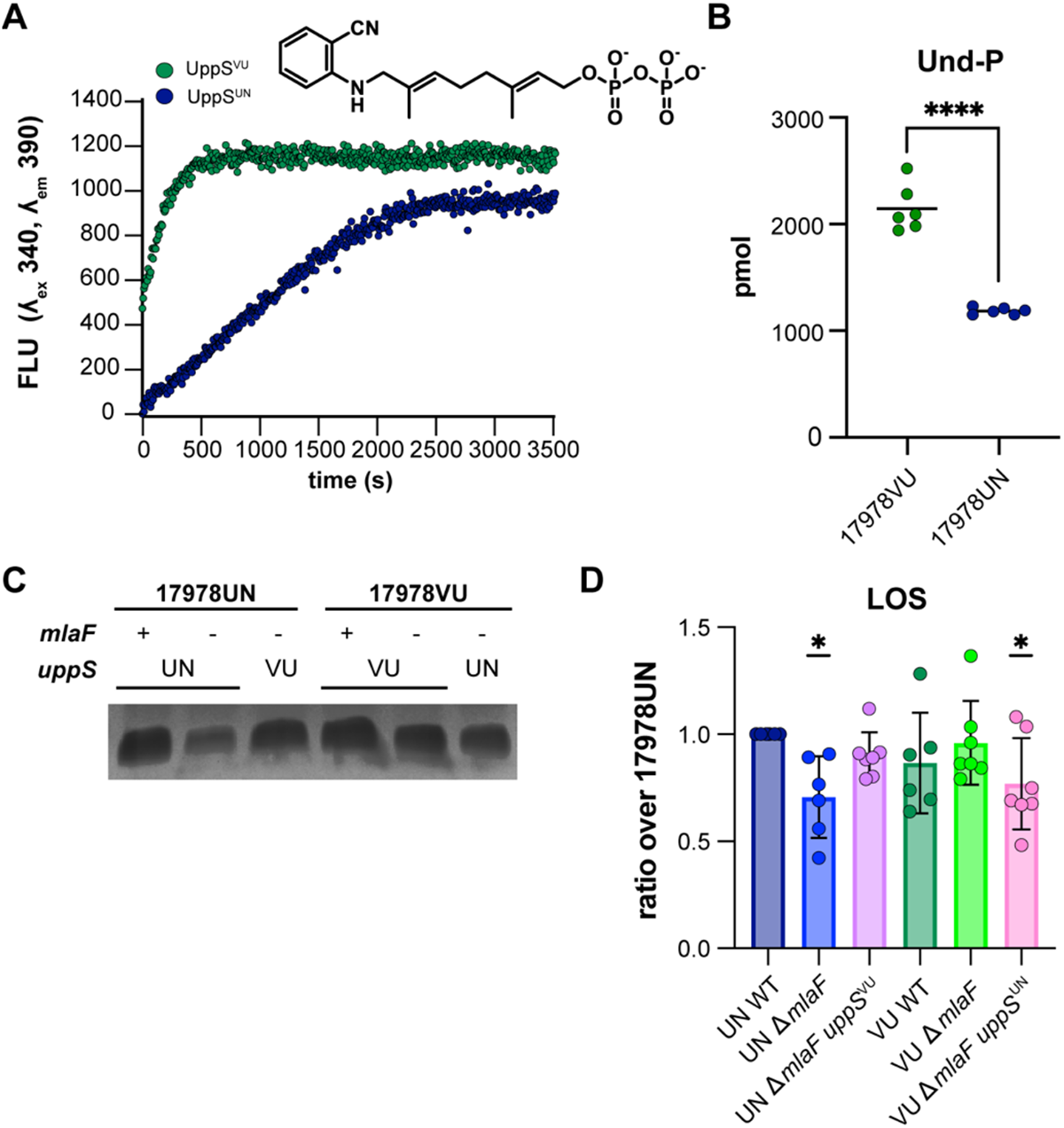
UppS^UN^ demonstrates decreased enzymatic rate and results in decreased cellular Und-P than UppS^VU^. **(A)** UppS activity with 2CNA-GPP, a fluorescent analogue of the intrinsic substrate FPP. *A. baumanni* UppS from strains 17978VU (green) and 17978UN (blue) and were tested with the 2CNA-GPP substrate analogue and fluorescence increase upon elongation was monitored at an excitation wavelength of 340 nm and emission of 390 nm. Data are representative of 3 experiments. **(B)** Mass spectrometry quantitation of undecaprenyl-phosphate in ATCC 17978VU and 17978UN WT strains. Significance is by an unpaired *t*-test; **** *p* < 0.0001. **(C)** Silver stain of proteinase K-treated cell lysates to stain for LOS levels. Image is representative of 7 biological replicates. **(D)** Densitometric quantification of LOS bands as a ratio over 17978UN WT. n=6-7, data are means ± SEM, significance is by one-sample *t*-test compared to 1; * *p* < 0.05.

Und-P has an established role in capsule, peptidoglycan, and glycoprotein biosynthesis; therefore, we hypothesized that capsule content or the cell wall may be altered by the reduced activity of UppS^UN^. However, Maneval staining of capsule and 3-[(7-Nitro-2,1,3-benzoxadiazol-4-yl)amino]-D-alanine hydrochloride (NADA) peptidoglycan stains showed no observable differences among strains (Fig. S2C-D). In other Gram-negative organisms, Und-P has an established role in the production of LPS through as the lipid carrier of O-antigen precursors (11). There is no known role for Und-P in the biosynthesis of LOS that lacks the O-antigen. Nevertheless, we previously found that the *A. baumannii* 17978UN Δ*mlaF* mutant contained significantly reduced levels of LOS compared to the wildtype strain (20). We therefore hypothesized that if Und-P plays a role in LOS biosynthesis, there would be reduced levels of LOS in the 17978UN Δ*mlaF* and 17978VU *mlaF uppS*^UN^ strains. Indeed, when LOS was quantified by silver staining of proteinase K-treated cell lysates, Δ*mlaF* strains with *uppS*^UN^ had reduced LOS abundance (Fig. 5C-D, Fig. S2E). The WT strains and the Δ*mlaF* strains with *uppS*^VU^ showed no reduction in LOS (Fig. 5C-D, Fig. S2E). While further experiments would be required to confirm a role for UppS and Und-PP in LOS production, our data suggests that an aberrant UppS results in reduced LOS. Together, this data suggests that UppS^UN^ results in reduced LOS abundance in an Δ*mlaF* mutant that confers a defective cellular envelope barrier function.

### Mla and uppS synergy influence clinically relevant phenotypes

Previous reports showed that *A. baumannii mla* mutants have increased sensitivity to antimicrobials such as gentamicin, novobiocin, rifampicin, meropenem, and the superoxide donor paraquat (20, 23, 49). We hypothesized that in a Δ*mlaF* mutant background, the *uppS*^UN^ allele would result in higher sensitivity to antimicrobials than the *uppS*^VU^ allele. In a disc diffusion assay, the presence of *uppS*^UN^ in a Δ*mlaF* mutant resulted in increased sensitivity to multiple antimicrobials, including first line antibiotics such as meropenem and imipenem, regardless of the ATCC 17978 background (Fig. 6A-D). These results show that UppS synergizes with the Mla system to promote *A. baumannii* antimicrobial resistance. As expected, the *ispB* suppressor only restored resistance to strains containing the *uppS*^UN^ allele. These data are consistent with the model that the isoprenoid biosynthetic pathway contributes to antimicrobial resistance in the Δ*mlaF* mutant by altering flux to Und-P via UppS. We further hypothesized that the *uppS*^UN^ allele would confer increased sensitivity to lysozyme, a host-produced enzyme that cleaves the peptidoglycan cell wall, compared to the *uppS*^VU^ allele in a Δ*mlaF* mutant (60). During growth in LB with 1 mg/mL lysozyme, the 17978UN Δ*mlaF* exhibited severe susceptibility to lysozyme compared to the UN WT strain (Fig. 6E, Fig. S3E). Resistance was restored to 17978UN Δ*mlaF* by the *uppS*^VU^ allele. Similarly, the *uppS*^UN^ allele conferred greater susceptibility to 17978VU Δ*mlaF* (Fig. 6E, Fig. S3E). Together, these data show that the *uppS*^VU^ allele helps to resist multiple antimicrobial stresses in a Δ*mlaF* mutant.

**Figure 6.**
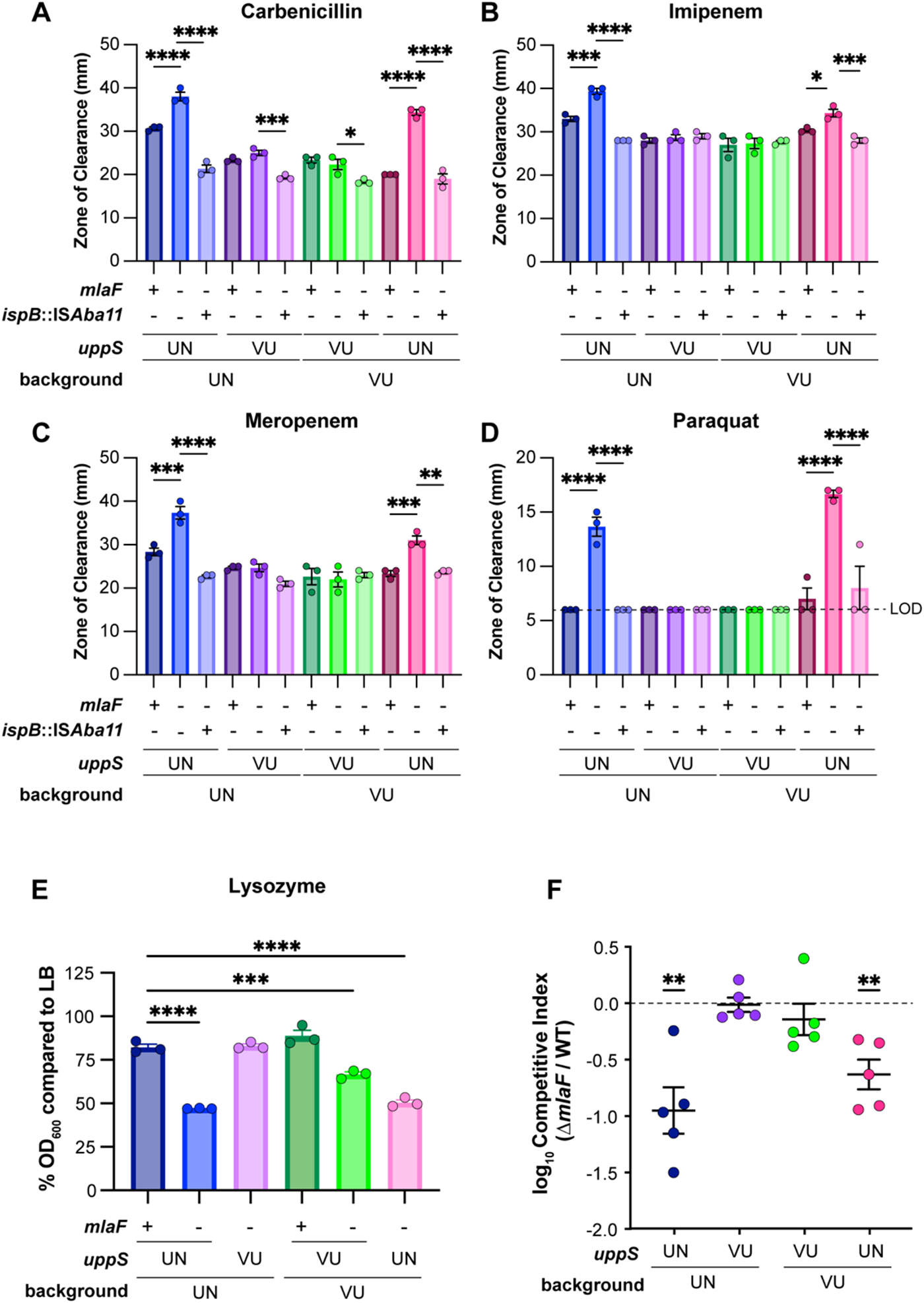
UppS^UN^ confers greater susceptibility to antimicrobials and decreased virulence in a murine model of pneumonia in an Δ*mlaF* background. **(A to D)** Antimicrobial susceptibility of VU and UN WT and mutant strains was determined by a disk-diffusion assay measuring the zone of clearance. Data are representative of 3 experiments. n=3, data are mean ± SEM. Significance is by one-way ANOVA with Tukey’s multiple comparisons. Limit of detection (LOD) = 6 mm. **(E)** Lysozyme susceptibility of VU and UN WT and mutant strains at 12 h during growth with a final concentration of 1 mg/mL lysozyme. n=3, data are means ± SEM. Significance is by one-way ANOVA with Tukey’s multiple comparisons. **(F)** VU and UN WT and Δ*mlaF* strains with the intrinsic or opposite *uppS* allele were used to intranasally infect C57BL/6 mice in a 1:1 WT:mutant ratio. After 48 h post infection, the lungs were harvested and bacterial burdens were enumerated. n=5, normality of log_10_-transformed competitive index data was determined by the Kolmogorov-Smirnov test and significance competitive index was determined by a one sample *t* test compared to 0. * *p* < 0.05, ** *p* <0.01, *** *p* < 0.001, **** *p* < 0.0001.

We previously reported that the Δ*mlaF* mutant in the ATCC 17978UN background had a virulence defect in a murine model of pneumonia (20). We hypothesized that the Δ*mlaF uppS*^UN^ strain would have a greater virulence defect than the Δ*mlaF uppS*^VU^ strain. Therefore, we compared bacterial burdens in a competitive infection with WT 17978VU and 17978UN strains and Δ*mlaF* mutants where the mutant expressed either the intrinsic or opposite *uppS* allele. At 40 h post infection, the Δ*mlaF* mutants containing the *uppS*^UN^ allele had a significant defect in virulence compared to WT regardless of strain background (Fig. 6F, Fig. S3A-D). The Δ*mlaF* mutant strains with *uppS*^VU^ showed no virulence defect compared to WT (Fig. 6F, Fig. S3A-D). This demonstrates that the virulence defect observed in Δ*mlaF* is due to synergy with the *uppS*^UN^ allele. Together, these data suggest that the Mla system and Und-P levels synergize to promote antimicrobial resistance and virulence in *A. baumannii*.

## Discussion

The *A. baumannii* cell envelope is the first line of defense against antibiotic stress and the host. Therefore, it is critical to understand the biological processes that uphold this barrier. The Mla system is an important factor in maintaining outer membrane lipid asymmetry to promote envelope function in Gram-negative bacteria. Multiple reports have demonstrated envelope defects in the absence of the Mla system in *A. baumannii* (23, 49, 61). Here, we found that the deletion of *mlaF* in two closely related *A. baumannii* type strains results in differential phenotypes. Genetic dissection uncovered synergy between the Und-P abundance and the Mla system in *A. baumannii*.

*A. baumannii* ATCC 17978 is a commonly used type strain. We recently discovered that ATCC 17978 was a mixed culture of two closely related strains that differed by the accessory locus *Aba*L44 and multiple SNPs (52). Depending on the time of order, the ratio of 17978UN to 17978VU isolates received from ATCC varied. In 2009, 4/6 isolates screened were 17978UN and 2/6 were 17978VU; by contrast, an ATCC order from 2021 yielded 35/36 as 17978UN and 1/36 as 17978VU (52). Differences in Δ*mlaF* phenotype of ATCC 17978UN and 17978VU presented the opportunity to use these variants as a genetic tool to investigate synthetic gene pairs. Ultimately, *uppS* was shown to synergize with Δ*mlaF* in resistance to membrane stresses and antibiotics, OM permeability, and virulence in a mouse lung infection. This result was surprising as a previous report identified SNPs in *obgE,* which encodes an essential GTPase involved in the stringent response, as the critical determinant of two isogenic ATCC 17978UN variants’ differences in growth and the stringent response (49). While we did not observe overt differences in growth based on the *obgE* allele, we speculate that is likely due to differences in growth conditions. Additionally, our results show that *obgE* is not synergistic with the Mla system for membrane stress resistance. Together, these findings exemplify that closely related strains can be leveraged to uncover underlying genetic synergy in *A. baumannii*.

As an essential glycan carrier, Und-P plays a role in the biosynthesis of multiple bacterial cell envelope components including peptidoglycan, glycoproteins, LPS O-antigen, lipid A, and capsular polysaccharides (51, 62, 63). However, *A. baumannii* does not synthesize the LPS O-antigen and does not encode LpxT that uses Und-PP as a phosphate donor for lipid A. Imaging of isogenic strains with UppS variants showed no overt difference in capsule or peptidoglycan (Fig. S2C-D). UppS^UN^ has a decreased enzymatic rate and confers increased membrane sensitivity in an Δ*mlaF* background. However, there were no defects in the wild-type background strains for membrane stress sensitivity, suggesting that the Δ*mlaF* defects synergize with reduced Und-P to result in a weakened cell envelope. ABUW5075, a clinical multidrug resistant isolate of *A. baumannii*, possess UppS^VU^ but demonstrated increased sensitivity to SDS/EDTA in a Δ*mlaF* background (20). This suggests that genetic components other than *uppS* may be at play in upholding the cell envelope in the absence of the Mla system. We previously observed that *A. baumannii* 17978UN Δ*mlaF* had decreased LOS abundance (20), suggesting Und-P may be important for LOS abundance in *A. baumannii*. Here, we show that the reduction in LOS within 17978UN Δ*mlaF* is due to UppS^UN^ as an isogenic mutant with UppS^VU^ displays WT-like LOS levels. This suggests that Und-P has an uncharacterized role in LOS synthesis in *A. baumannii*. For example, a non-homologous enzyme may use Und-PP as a phosphate donor for lipid A similar to LpxT. Alternatively, Und-P may serve as a glycan carrier for LOS such as for the sugar molecules in the core oligosaccharides.

Suppressor mutants in genes encoding isoprenoid biosynthetic pathway enzymes restored membrane stress resistance to Δ*mlaF uppS*^UN^ strains. We found that the Δ*mlaF uppS*^UN^ phenotype is suppressed by mutations in isoprenoid biosynthetic genes *ispA* and *ispB*. The IS*Aba11* transposition to the 5’ untranslated region of *ispB* was also found in an extensively drug resistant clinical isolate of *A. baumannii*, demonstrating potential clinical implications for the suppressive mechanism (64). Here, we identified another suppressor in an isoprenoid biosynthetic gene encoding IspA^G223E^ that arose when constructing the 17978VU Δ*mlaF uppS*^UN^ strain. In *E. coli*, *ispA* null mutants had reduced isoprenoid quinone levels that was rescued by the overexpression of either *ispB* or *ispU* (*uppS*) (65, 66). This suggests that there may be low level functional redundancy between IspA, IspB, and UppS. We therefore hypothesize that IspA^G223E^ enhances low-level Und-PP synthesis performed by IspA. We observed both *ispA* and *ispB* suppressors restoring growth to strains containing *uppS*^UN^, suggesting that *ispA* and *ispB* may compensate for a reduced activity of UppS in the 17978UN background. Similarly, in *E. coli*, an aberrant UppS mutant resulted in growth defects that were rescued by mutants in the isoprenoid pathway (67). This suggests that there may be a conserved mechanism to promote the production of undecaprenyl species in the presence of a defective UppS.

The structure of UppS has been solved from multiple organisms, including *A. baumannii* (68–70). The substitution in UppS that differs between 17978VU (M78) and 17978UN (T78) is at the 78^th^ amino acid position, located on the α3 helix. The α3 helix not only borders the active site but surrounds the open cavity the product chain would occupy. Residues along the α3 helix are largely conserved (71). The methionine at this position on the α3 helix is conserved in prenyl-transferases from *E. coli*, *Saccharomyces cerevisiae*, *Mycobacterium tuberculosis*, *Arabidopsis thaliana*, and humans (72). This suggests that the methionine is important for the structure or function of prenyl-transferases. In the 17978UN UppS variant the methionine at position 78 is replaced with a threonine. While not highlighted as a critical residue, the surrounding hydrophobic residues near *E. coli* UppS M86 (M78 in *A. baumannii*) (residues 85, 88, and 89 in *E. coli*) are implicated in interactions with the substrate (72). Importantly, there were no major structural differences in UppS between the M78 and T78 variants by circular dichroism analysis (Fig. S2A-B). The substitution to a threonine in 17978UN may therefore perturb internal hydrophobic interactions between the α3 helix and substrate. We found that 17978UN UppS has a reduced enzymatic rate compared to 17978VU UppS. This suggests that the T78 encoded by ATCC 17978UN may perturb the crucial interactions with the substrate hydrophobic tail required for Und-PP production.

Isoprenoid biosynthesis is an essential process in bacteria. Additionally, isoprenoids have diverse functions in the cell ranging from metabolism to virulence at the host-pathogen interface. As such, proteins within isoprenoid biosynthesis have been considered as a drug target for novel therapeutics (73, 74). Furthermore, UppS specifically is a proposed drug target for *A. baumannii* (75). The synthetically sick Δ*mlaF* mutants in ATCC 17978UN suggest that targeting UppS in a combination therapy may represent an effective therapeutic strategy. The genetic synergy between the *A. baumannii* Mla system and UppS highlights the importance of cell envelope maintenance during environmental stress and host infection. The cell envelope, and more specifically the OM, is the largest barrier to effective antibiotic treatment. Our findings provide a deeper understanding into how *A. baumannii* maintains cell envelope integrity to survive in hostile environments.

## Methods

### Bacterial strains and growth

All bacterial strains and plasmids used in this study are listed in Tables S1 and S2, respectively. Strains were grown in LB or on LB plates with 1.5% w/v agar. Antibiotics were used at the following concentrations: carbenicillin, 75 mg/L; kanamycin, 40 mg/L; chloramphenicol, 15 mg/L. Overnight cultures were started in 3-5 mL LB, inoculated with a single colony, and incubated at 37°C while shaking at 180 rpm for 8-16 hours. Growth curves were conducted in 100 μL media in a flat bottom 96-well plate, inoculated with 1 μL overnight culture, and incubated at 37°C with shaking. Bacterial growth was monitored by optical density at 600 nm (OD_600_) in an EPOCH2 BioTek (Winooski, VT) plate reader. For assays on membrane stress, SDS/EDTA was included in the media at varying concentrations and the concentrations used are noted in each figure due to variability. For lysozyme susceptibility assays, lysozyme was included in the media at a final concentration of 1 mg/mL and the OD_600_ at 12 hours was used to calculate the %OD_600_ compared to the average of the OD_600_ at 12-hours in LB for each strain.

### Plasmid construction and allelic exchange mutants

All oligonucleotides used are listed in Table S3. DNA was amplified using 2X Phusion Master Mix (ThermoFisher, Waltham, MA), Q5 High Fidelity 2X Master Mix (New England Biolabs (NEB), Ipswich, MA) or GoTaq Green Master Mix (NEB, Ipswich, MA). The pFLP2 vector was used for all allelic exchange mutants. Using ATCC 17978VU or 17978UN as the template, 1000 bp upstream and downstream of the mutation of interest was amplified. For pFLP2-*obgE*, HN1 and HN2 were used. For pFLP2-*lptD*, HN23 and HN24 were used. For pFLP2-*uppS*, HN25 and HN26 were used. For *ispA*\* reconstruction, HN85 and HN86 were used with strain LP546 as the template. The PCR product was incorporated into a digested pFLP2 backbone using KpnI and BamHI restriction sites and HiFi ligation mix (NEB, Ipswich, MA). All restriction enzymes are from NEB (Ipswich, MA). *A. baumannii* was transformed through conjugation by triparental mating with *E. coli* strain HB101 containing pRK2013 as the helper strain. Merodiploids containing the integrated pFLP2 vector were screened on plates containing 10% sucrose and 75 mg/L carbenicillin and Suc^S^ and Carb^R^ colonies were selected. Merodiploids with the appropriate phenotype were screened by PCR to confirm plasmid incorporation at the correct site. Merodiploids were grown on LB agar, resuspended in LB, and plated to LB agar containing 10% sucrose to select for second crossover events, and the Suc^R^ strains were screened for Carb^S^. Genotypes were confirmed via Sanger Sequencing (UIC Genomics Research Core) and/or whole genome sequencing (SeqCoast Genomics, Portsmouth New Hampshire; SeqCenter, Pittsburg PA).

Strains containing the *ispB* suppressor mutation and *mlaF* knockout were generated using pLDP70 and pLDP8, respectively (20).

UppS expression plasmids for protein purification were generated using pET-15b digested with BamHI and NdeI. UppS was amplified from either 17978VU or 17978UN with HN94 and HN95 and ligated into the pET-15b backbone using a HiFi ligation. All plasmid sequences were confirmed with Sanger Sequencing by the UIC Genomics Research Core or whole plasmid sequencing by Primordium (Monrovia, CA).

### Serial Dilution Spotting Assays

Overnight cultures were 10-fold serially diluted in 96-well plates in 1X PBS to 10^-7^. Dilutions were spotted in 3 or 5 μL drops on LB agar and LB agar containing SDS/EDTA and incubated at 37°C overnight. Images of plates were taken on a BioTek (Winooski, VT) ChemiDoc MP imager.

### Antibiotic susceptibility assay

9.5 mL aliquots of soft agar containing 0.8% agar were inoculated with 270 μL overnight culture and plated to a prewarmed 15-cm LB agar plate. Once solidified, pre-loaded antibiotic discs (BD (Becton Dickinson), Franklin Lakes, NJ) were placed using the BD automatic disc dispenser onto the agar overlay and the plates were incubated at 37°C overnight. The following day, diameters of the zones of clearance were measured in mm.

### Ethidium bromide uptake assay

The EtBr uptake assay was adapted from previous studies (23, 57). Bacteria were grown in 3 mL LB to mid-log phase before centrifuging and normalizing to ∼1x10^10^ CFU in PBS. Normalized cells were plated to confirm equal CFU across strains. In 200 μL final volume, normalized cells were combined with 200 μM carbonyl cyanide 3-chlorophenylhydrazone (CCCP) and 1.2 μM EtBr. EtBr uptake was monitored in a black 96-well plate with reads every 15 s on a BioTek Synergy H1 (Winooski, VT) using excitation and emission wavelengths of 530 nm and 590 nm, respectively.

### Maneval stain for capsule

Strains were streaked to LB and incubated overnight at 37°C. A single colony was resuspended in 10 μL of 1% Congo Red and spread across a microscope slide. Slides were flooded with Maneval Stain (acid fuchsin, glacial acetic acid, iron(III) chloride)(76) for 5 minutes before gentle washing and imaging at 60X on a Keyence BZ-X Inverted Fluorescence microscope (Itasca, IL).

### NADA-green peptidoglycan stain

3-[(7-Nitro-2,1,3-benzoxadiazol-4-yl)amino]-D-alanine hydrochloride (NADA) was used as previously described (77). Cultures were grown overnight at 37°C. The following morning, cells were subcultured into fresh media and grown to mid-log phase with an OD_600_ of 0.5. 1 mL of OD_600_ 0.5 cells were pelleted, washed in LB, and resuspended in 1 mL fresh LB. NADA-green (ThermoFisher, Waltham, MA) was added to the resuspended cells to a final concentration of 30 μM and incubated for 30 minutes at 37°C. Cells were then fixed in 1.6% paraformaldehyde and plated on an agar pad. Imaging was performed at 60X using a Keyence BZ-X Inverted Fluorescence microscope (Itasca, IL).

### UppS purification

UppS^VU^ and UppS^UN^ were purified from BL21 ArcticExpress DE3 RIL cells (VWR, Radnor, PA). Cells expressing pET-15b-UppS^VU^ or pET-15b-UppS^UN^ were grown in 10 mL LB containing carbenicillin overnight at 37°C. The following day, cells were subcultured into 1 L LB containing carbenicillin in a 2.8 L Fernbach flask and grown to mid-log phase at 37°C. At OD_600_ of 0.6, IPTG was added to a final concentration of 0.5 mM to induce protein expression. Cells were incubated overnight while shaking (180 rpm) at 25°C. The following day cells were centrifuged at 2,000 x *g* for 10 minutes and the cell pellet was lysed using 8 mL of B-PER Bacterial Protein Extraction Reagent (Thermo Scientific, Waltham, MA) per gram of pellet with gentle shaking for 1 hour. The lysate was pelleted by centrifugation at 4,300 x *g* for 5 minutes. The soluble fraction in the supernatant was mixed with an equal part lysis buffer (50 mM NaH_2_PO_4_, pH 8.0, 300 mM NaCl, 10 mM imidazole) and was added to 8 mL Ni-NTA resin (Qiagen, Hilden, Germany) preequilibrated with lysis buffer and rocked for 1 hour at 4°C. Protein-bound resin was then applied to a 10-mL chromatography column (BioRad, Hercules, CA) and washed twice with 10 mL wash buffer (50 mM NaH_2_PO_4_, pH 8.0, 300 mM NaCl, 20 mM imidazole). Protein was eluted using elution buffer (50 mM NaH_2_PO_4_, pH 8.0, 300 mM NaCl, 250 mM imidazole) in serial 2 mL elution volumes. Samples were separated by SDS-PAGE and stained with SimplyBlue Safe Stain (Invitrogen, Waltham, MA) to verify protein purification. Protein was desalted using PD-10 desalting columns (Cytiva, Marlborough, MA) into circular dichroism and UppS assay buffers (see below).

### Circular dichroism

UppS^VU^ and UppS^UN^ were prepared to 0.1 mg/mL in circular dichroism (CD) buffer (10 mM sodium phosphate buffer, pH 8.0, with 10% glycerol). CD spectra were collected by the UIC BioPhysics Core from a 0.1 mg/mL sample in a 1 mm cuvette from 190 nm to 240 nm on a Jasco 815. Protein secondary structure was determined by the software CDSSTR (78, 79).

### Undecaprenyl-phosphate quantification

5 mL cultures of 17978UN and 17978VU were grown in LB to mid-log phase with a final OD_600_ of 0.5. Cultures were pelleted and frozen at -80 for further processing. Pellets were then subjected to Bligh and Dyer extraction and dried overnight. The crude cell lysate was then resuspended in 200 mL of *n*-propanol and 0.1 % ammonium hydroxide (1:3)(80). After sonicating the cell suspension using water bath, 5 mL of sample was injected into C18 column and analyzed for C55 bactoprenol phosphate (BP) m/z ratio of 845.7 using LC-MS. Area under the curve of each BP peak of all samples were recorded and used to calculate BP (pmol) using a standard curve generated from known BP concentrations.

### UppS microplate enzyme assays

2CNA-GPP was prepared as described previously (58, 67). In each well of a black-walled 96 well plate, reactions were prepared with 2.5 mM 2CNA-GPP, 0.5 mM MgCl2, 5 mM KCl, 0.1 % Triton-X-100, and 100 nM recombinant UppS from 17978UN or 17978VU. The plate was incubated in the plate reader at 30° C for five minutes then the reaction was initiated with the addition of 1 mM IPP (final concentration). Fluorescence was monitored at 30°C over one hour at an excitation wavelength of 340 nm and emission wavelength of 390 nm. The reaction rate was determined based on the linear fluorescence increase over the first 8 minutes for both strains.

### LOS silver stain and densitometry

Samples were prepared similarly to previous descriptions (81, 82). Three-mL overnight cultures were diluted 1:100 in 5 mL fresh LB and grown to mid-log with an OD_600_ of 0.5. Two 1-mL aliquots were harvested. One aliquot was boiled at 85°C for 15 min and subjected to a Pierce 660 (Thermo Fisher, Waltham, MA) total protein quantification following manufacturer instructions. The second aliquot was pelleted and resuspended in 1X Novex SDS sample buffer (Invitrogen, Waltham, MA) so that the final total protein concentration was 6 mg/ml, which was determined to be in the linear range of the LOS densitometry analysis by a dilution series experiment. Samples were lysed via boiling for 15 min at 85°C and treated with proteinase K to a final concentration of 0.16 μg/μl. Gels were loaded with samples containing 10 μg total protein before electrophoresis on a 16.5% Tris Tricine gel (Bio-Rad, Hercules, CA). Gels were stained with the Pierce Silver Stain kit (Thermo Fisher, Waltham, MA) following manufacturer instructions. Images were captured on a BioTek (Winooski, VT) ChemiDoc MP imager.

Densitometry analysis was performed using ImageJ. Images were converted to an 8-bit image and the background was subtracted using a 50-pixel rolling ball radius. Using the gel analysis tool, LOS bands were selected and peaks were quantified. Data are presented as a ratio of the value of 17978UN WT for each independent gel.

### Murine model of A. baumannii lung infection

Six-week-old female C57BL/6 mice were purchased from Jackson Laboratory. Mice were housed in a temperature-controlled environment with 12-h light/dark cycles and food and water were provided as needed and were acclimated to the facility to 1 week prior to infection. Mice were anesthetized with ketamine/xylazine and inoculated intranasally with 35 μL bacterial suspension 1:1 mixture *A. baumannii* ATCC 17978 WT and Δ*mlaF* mutant derivatives, containing approximately 3 × 10^8^ CFU as described previously (20). The inoculum dose was confirmed by serial dilution and plating on selective agar media. Mice were euthanized at 48 h post infection by CO_2_ asphyxiation and the lungs were excised aseptically. Tissues were homogenized and all samples were serially diluted and plated on LB and kanamycin selective agar plates for bacterial enumeration. The competitive index of mutant/WT strains were calculated by dividing the mutant CFU ratio (CFU_output_/CFU_input_) by the WT CFU ratio. All animal care protocols were approved by the University of Illinois Chicago Institutional Animal Care and Use Committee (IACUC; protocol number 20-165) in accordance with the Animal Care Policies of UIC, the Animal Welfare Act, the National Institutes of Health, and the American Veterinary Medical Association (AVMA). Animals were humanely euthanized consistent with the AVMA guidelines.

### Variant protein tree construction – genome filtering and de-duplication

To acquire genomes for analysis, 7431 summaries of *Acinetobacter baumannii* reference assemblies (assembly accessions starting with “GCF_”) were downloaded from NCBI datasets using the datasets command line interface tool. Genomes with contig N50 scores in the bottom 20% were removed from the assemblies, resulting in a total of 5945 genomes. Since there were many closely related genomes, a deduplication process was carried out based on Average Nucleotide Identity (ANI) values estimated using Mash distances generated by Mash version 2.34 (83). Each genome was represented by a compressed min-Hash sketch, and pairwise distances between sketches were used to construct a standard distance matrix. To perform the deduplication, a custom Python script iterated through each genome in the distance matrix, comparing its distances to other genomes. A chosen threshold Mash distance (t_dist_) of 0.006 was used. If the pairwise distance between two genomes was less than t_dist_, the genome with the lower N50 score was discarded. This step was performed recursively until all pairwise distances were greater than t_dist_, ensuring that only unique genomes remained in the dataset. Ten assemblies were identified as outliers due to their high average Mash distance to all other genomes and were excluded. Scripts for downloading genomes, creating Mash sketches and distance matrices, and dereplicating genomes and removing outliers are available at https://github.com/JonWinkelman/genome_deduplication.

### Variant protein tree construction – identification of orthologs in A. baumannii

We utilized OrthoFinder (54, 55) version 2.5.4 with default settings to determine orthologous relationships between genes in *A. baumannii* proteomes. A total of 233 proteomes from filtered and de-duplicated *Acinetobacter* genomes were included in the analysis. These genomes include three outgroups*, A. baylyi*, *A. gyllenbergii* and *A. colistiniresistens* that were used to root the species tree. OrthoFinder computed hierarchical orthologous groups (HOGs) for each internal node in the species tree. These HOGs consist of proteins descended from a single gene in the ancestral species corresponding to the respective internal node. For this study, we focused on analyzing HOGs associated with the species tree node representing the last common ancestor of all *A. baumannii*. OrthoFinder computes HOGs for each internal node of the species tree. Each HOG contains all genes that descended from a single gene in ancestral species represented by the internal node. In this analysis we analyzed HOGs for the species tree node representing the last common ancestor of all *A. baumannii.* A custom python dash application was used to explore results from OrthoFinder, and iTOL was used to annotate trees for figures (84). The proteomes and alleles depicted are listed in Table S4.

### Variant protein tree construction – Manual identification of orthologs in A. baumannii

When working with three specific genes, OrthoFinder search for orthologs didn’t yield results across all species. In cases where OrthoFinder didn’t locate an ortholog, we adopted an alternative approach. Specifically, we investigated the presence or absence of neighboring genes. For instance, if an ortholog for the *Acinetobacter baumannii* 17978-mff gene *ACX60_11495* was detected in a strain, orthologs to all other genes in its operon were found. Conversely, when this ortholog was absent, none of the operon’s other genes were found, suggesting that this gene and its operon were indeed absent from the genome.

In another instance involving the *Acinetobacter baumannii* 17978-mff gene *ACX60_05080*, OrthoFinder failed to identify orthologs in two strains. In this case, we observed adjacent orthologs and identified a potential ortholog with an unassigned HOG and more than 95% sequence identity to ACX60_05080. These findings strongly suggested that this identified potential ortholog was indeed an orthologous gene. Jupyter notebooks for downloading genomes, creating Mash sketches and distance matrices, dereplicating genomes, removing outliers, processing OrthoFinder results and creating figures are available at https://github.com/JonWinkelman/Palmer_baumannii_UNvVUprevalence.git.

## Data availability

Whole genome sequencing data is available in the National Center for Biotechnology Information (NCBI) sequence read archive (SRA) under BioProject: PRJNA1020123 and PRJNA656143 (20).

## Acknowledgements

This work was supported by startup funds from the University of Illinois Chicago and National Institutes of Health Awards R00HL143440 to L.D.P. and R01GM123251 to J.M.T. We thank the UIC Biophysics Core and the Center for Structural Biology for assisting with the circular dichroism experiment and analysis. We thank members of the Palmer laboratory for critical reading of the manuscript and Zachery Lonergan for review of the manuscript. The authors declare no conflict of interest regarding the content of this manuscript.

## Author contributions

H.R.N. – project conceptualization, methodology, data collection, data analysis, and writing. S.K. – data collection and writing. X.R. – data collection and writing. J.D.W – bioinformatics, data analysis, and writing. J.R.T. – data collection, data analysis, writing, funding, and resources. L.D.P. – project conceptualization, methodology, funding, resources, data analysis, and writing.

## References

1. Han L, Gao Y, Liu Y, Yao S, Zhong S, Zhang S, Wang J, Mi P, Wen Y, Ouyang Z, Zhang J, Al-Shamiri MM, Li P, Han S. 2022. An outer membrane protein YiaD contributes to adaptive resistance of meropenem in *Acinetobacter baumannii*. Microbiol Spectr 10:e0017322.

2. Trebosc V, Gartenmann S, Tötzl M, Lucchini V, Schellhorn B, Pieren M, Lociuro S, Gitzinger M, Tigges M, Bumann D, Kemmer C. 2019. Dissecting colistin resistance mechanisms in extensively drug-resistant *Acinetobacter baumannii* clinical isolates. mBio 10:e01083–19.

3. 3. World Health Organization. 2019. No time to wait: securing the future from drug-resistant infections. Report to the Secretary-General of the United Nations. World Health Organization and Interagency Coordination Group on Antimicrobial Resistance.

4. Weiner LM, Webb AK, Limbago B, Dudeck MA, Patel J, Kallen AJ, Edwards JR, Sievert DM. 2016. Antimicrobial-resistant pathogens associated with healthcare-associated infections: summary of data reported to the National Healthcare Safety Network at the Centers for Disease Control and Prevention, 2011–2014. Infect Control Hosp Epidemiol 37:1288–1301.

5. Tacconelli E, Carrara E, Savoldi A, Harbarth S, Mendelson M, Monnet DL, Pulcini C, Kahlmeter G, Kluytmans J, Carmeli Y, Ouellette M, Outterson K, Patel J, Cavaleri M, Cox EM, Houchens CR, Grayson ML, Hansen P, Singh N, Theuretzbacher U, Magrini N, Aboderin AO, Al-Abri SS, Awang Jalil N, Benzonana N, Bhattacharya S, Brink AJ, Burkert FR, Cars O, Cornaglia G, Dyar OJ, Friedrich AW, Gales AC, Gandra S, Giske CG, Goff DA, Goossens H, Gottlieb T, Guzman Blanco M, Hryniewicz W, Kattula D, Jinks T, Kanj SS, Kerr L, Kieny M-P, Kim YS, Kozlov RS, Labarca J, Laxminarayan R, Leder K, Leibovici L, Levy-Hara G, Littman J, Malhotra-Kumar S, Manchanda V, Moja L, Ndoye B, Pan A, Paterson DL, Paul M, Qiu H, Ramon-Pardo P, Rodríguez-Baño J, Sanguinetti M, Sengupta S, Sharland M, Si-Mehand M, Silver LL, Song W, Steinbakk M, Thomsen J, Thwaites GE, van der Meer JW, Van Kinh N, Vega S, Villegas MV, Wechsler-Fördös A, Wertheim HFL, Wesangula E, Woodford N, Yilmaz FO, Zorzet A. 2018. Discovery, research, and development of new antibiotics: the WHO priority list of antibiotic-resistant bacteria and tuberculosis. Lancet Infect Dis 18:318–327.

6. Geisinger E, Huo W, Hernandez-Bird J, Isberg RR. 2019. *Acinetobacter baumannii*: envelope determinants that control drug resistance, virulence, and surface variability. Annu Rev Microbiol 73:481–506.

7. Islam N, Kazi MI, Kang KN, Biboy J, Gray J, Ahmed F, Schargel RD, Boutte CC, Dörr T, Vollmer W, Boll JM. 2022. Peptidoglycan recycling promotes outer membrane integrity and carbapenem tolerance in *Acinetobacter baumannii*. mBio 13:e01001–22.

8. Silhavy TJ, Kahne D, Walker S. 2010. The Bacterial cell envelope. Cold Spring Harb Perspect Biol 2:a000414.

9. Rojas ER, Billings G, Odermatt PD, Auer GK, Zhu L, Miguel A, Chang F, Weibel DB, Theriot JA, Huang KC. 2018. The outer membrane is an essential load-bearing element in Gram-negative bacteria. 7715. Nature 559:617–621.

10. Moffatt JH, Harper M, Harrison P, Hale JDF, Vinogradov E, Seemann T, Henry R, Crane B, St. Michael F, Cox AD, Adler B, Nation RL, Li J, Boyce JD. 2010. Colistin resistance in *Acinetobacter baumannii* is mediated by complete loss of lipopolysaccharide production. Antimicrob Agents Chemother 54:4971–4977.

11. Simpson BW, Trent MS. 2019. Pushing the envelope: LPS modifications and their consequences. 7. Nat Rev Microbiol 17:403–416.

12. Simpson BW, Nieckarz M, Pinedo V, McLean AB, Cava F, Trent MS. 2021. *Acinetobacter baumannii* can survive with an outer membrane lacking lipooligosaccharide due to structural support from elongasome peptidoglycan synthesis. mBio 12:e03099–21.

13. Nikaido H. 2003. Molecular basis of bacterial outer membrane permeability revisited. Microbiol Mol Biol Rev 67:593–656.

14. Malinverni JC, Silhavy TJ. 2009. An ABC transport system that maintains lipid asymmetry in the Gram-negative outer membrane. Proc Natl Acad Sci 106:8009–8014.

15. Ekiert DC, Bhabha G, Isom GL, Greenan G, Ovchinnikov S, Henderson IR, Cox JS, Vale RD. 2017. Architectures of lipid transport systems for the bacterial outer membrane. Cell 169:273–285.e17.

16. Chong Z-S, Woo W-F, Chng S-S. 2015. Osmoporin OmpC forms a complex with MlaA to maintain outer membrane lipid asymmetry in *Escherichia coli*. Mol Microbiol 98:1133– 1146.

17. Thong S, Ercan B, Torta F, Fong ZY, Wong HYA, Wenk MR, Chng S-S. 2016. Defining key roles for auxiliary proteins in an ABC transporter that maintains bacterial outer membrane lipid asymmetry. eLife 5:e19042.

18. Ercan B, Low W-Y, Liu X, Chng S-S. 2019. Characterization of interactions and phospholipid transfer between substrate binding proteins of the OmpC-Mla System. Biochemistry 58:114–119.

19. Sutterlin HA, Shi H, May KL, Miguel A, Khare S, Huang KC, Silhavy TJ. 2016. Disruption of lipid homeostasis in the Gram-negative cell envelope activates a novel cell death pathway. Proc Natl Acad Sci 113:E1565–E1574.

20. Palmer LD, Minor KE, Mettlach JA, Rivera ES, Boyd KL, Caprioli RM, Spraggins JM, Dalebroux ZD, Skaar EP. 2020. Modulating isoprenoid biosynthesis increases lipooligosaccharides and restores *Acinetobacter baumannii* resistance to host and antibiotic stress. Cell Rep 32:108129.

21. MacRae MR, Puvanendran D, Haase MAB, Coudray N, Kolich L, Lam C, Baek M, Bhabha G, Ekiert DC. 2023. Protein-protein interactions in the Mla lipid transport system probed by computational structure prediction and deep mutational scanning. J Biol Chem 104744.

22. Isom GL, Davies NJ, Chong Z-S, Bryant JA, Jamshad M, Sharif M, Cunningham AF, Knowles TJ, Chng S-S, Cole JA, Henderson IR. 2017. MCE domain proteins: conserved inner membrane lipid-binding proteins required for outer membrane homeostasis. 1. Sci Rep 7:8608.

23. Kamischke C, Fan J, Bergeron J, Kulasekara HD, Dalebroux ZD, Burrell A, Kollman JM, Miller SI. 2019. The *Acinetobacter baumannii* Mla system and glycerophospholipid transport to the outer membrane. eLife 8:e40171.

24. de Jonge EF, Vogrinec L, van Boxtel R, Tommassen J. 2022. Inactivation of the Mla system and outer-membrane phospholipase A results in disrupted outer-membrane lipid asymmetry and hypervesiculation in *Bordetella pertussis*. Curr Res Microb Sci 3:100172.

25. Bernier SP, Son S, Surette MG. 2018. The Mla pathway plays an essential role in the intrinsic resistance of *Burkholderia cepacia* complex species to antimicrobials and host innate components. J Bacteriol 200:e00156–18.

26. Suzuki T, Murai T, Fukuda I, Tobe T, Yoshikawa M, Sasakawa C. 1994. Identification and characterization of a chromosomal virulence gene, *vacJ*, required for intercellular spreading of *Shigella flexneri*. Mol Microbiol 11:31–41.

27. Hong M, Gleason Y, Wyckoff EE, Payne SM. 1998. Identification of two *Shigella flexneri* chromosomal loci involved in intercellular spreading. Infect Immun 66:4700–4710.

28. Carpenter CD, Cooley BJ, Needham BD, Fisher CR, Trent MS, Gordon V, Paynea SM. 2014. The Vps/VacJ ABC transporter is required for intercellular spread of *Shigella flexneri*. Infect Immun 82:660–669.

29. Cuccui J, Easton A, Chu KK, Bancroft GJ, Oyston PCF, Titball RW, Wren BW. 2007. Development of signature-tagged mutagenesis in *Burkholderia pseudomallei* to identify genes important in survival and pathogenesis. Infect Immun 75:1186–1195.

30. Munguia J, LaRock DL, Tsunemoto H, Olson J, Cornax I, Pogliano J, Nizet V. 2017. The Mla pathway is critical for *Pseudomonas aeruginosa* resistance to outer membrane permeabilization and host innate immune clearance. J Mol Med 95:1127–1136.

31. Nasu H, Shirakawa R, Furuta K, Kaito C. 2022. Knockout of *mlaA* increases *Escherichia coli* virulence in a silkworm infection model. PLoS ONE 17:e0270166.

32. Baarda BI, Zielke RA, Van AL, Jerse AE, Sikora AE. 2019. *Neisseria gonorrhoeae* MlaA influences gonococcal virulence and membrane vesicle production. PLOS Pathog 15:e1007385.

33. Abellon-Ruiz J. 2022. Forward or backward, that is the question: phospholipid trafficking by the Mla system. Emerg Top Life Sci 7:125–135.

34. Powers MJ, Trent MS. 2019. Intermembrane transport: glycerophospholipid homeostasis of the Gram-negative cell envelope. Proc Natl Acad Sci 116:17147–17155.

35. Henderson JC, Zimmerman SM, Crofts AA, Boll JM, Kuhns LG, Herrera CM, Trent MS. 2016. The power of asymmetry: architecture and assembly of the Gram-negative outer membrane lipid bilayer. Annu Rev Microbiol 70:255–278.

36. Nagy E, Losick R, Kahnea D. 2019. Robust suppression of lipopolysaccharide deficiency in *Acinetobacter baumannii* by growth in minimal medium. J Bacteriol 201.

37. Powers MJ, Trent MS. 2018. Phospholipid retention in the absence of asymmetry strengthens the outer membrane permeability barrier to last-resort antibiotics. Proc Natl Acad Sci 115:E8518–E8527.

38. Low W-Y, Thong S, Chng S-S. 2021. ATP disrupts lipid-binding equilibrium to drive retrograde transport critical for bacterial outer membrane asymmetry. Proc Natl Acad Sci 118:e2110055118.

39. Awai K, Xu C, Tamot B, Benning C. 2006. A phosphatidic acid-binding protein of the chloroplast inner envelope membrane involved in lipid trafficking. Proc Natl Acad Sci 103:10817–10822.

40. Abellón-Ruiz J, Kaptan SS, Baslé A, Claudi B, Bumann D, Kleinekathöfer U, van den Berg B. 2017. Structural basis for maintenance of bacterial outer membrane lipid asymmetry. Nat Microbiol 2:1616–1623.

41. Tang X, Chang S, Qiao W, Luo Q, Chen Y, Jia Z, Coleman J, Zhang K, Wang T, Zhang Z, Zhang C, Zhu X, Wei X, Dong C, Zhang X, Dong H. 2021. Structural insights into outer membrane asymmetry maintenance in Gram-negative bacteria by MlaFEDB. Nat Struct Mol Biol 28:81–91.

42. Yeow J, Tan KW, Holdbrook DA, Chong Z-S, Marzinek JK, Bond PJ, Chng S-S. 2018. The architecture of the OmpC–MlaA complex sheds light on the maintenance of outer membrane lipid asymmetry in *Escherichia coli*. J Biol Chem 293:11325–11340.

43. Douglass MV, McLean AB, Trent MS. 2022. Absence of YhdP, TamB, and YdbH leads to defects in glycerophospholipid transport and cell morphology in Gram-negative bacteria. PLOS Genet 18:e1010096.

44. Ruiz N, Davis RM, Kumar S. 2021. YhdP, TamB, and YdbH are redundant but essential for growth and lipid homeostasis of the gram-negative outer membrane. mBio 12:e0271421.

45. Hughes GW, Hall SCL, Laxton CS, Sridhar P, Mahadi AH, Hatton C, Piggot TJ, Wotherspoon PJ, Leney AC, Ward DG, Jamshad M, Spana V, Cadby IT, Harding C, Isom GL, Bryant JA, Parr RJ, Yakub Y, Jeeves M, Huber D, Henderson IR, Clifton LA, Lovering AL, Knowles TJ. 2019. Evidence for phospholipid export from the bacterial inner membrane by the Mla ABC transport system. 10. Nat Microbiol 4:1692–1705.

46. Mann D, Fan J, Somboon K, Farrell DP, Muenks A, Tzokov SB, DiMaio F, Khalid S, Miller SI, Bergeron JRC. 2021. Structure and lipid dynamics in the maintenance of lipid asymmetry inner membrane complex of *A. baumannii*. Commun Biol 4:817.

47. Coudray N, Isom GL, MacRae MR, Saiduddin MN, Bhabha G, Ekiert DC. 2020. Structure of bacterial phospholipid transporter MlaFEDB with substrate bound. eLife 9:e62518.

48. Chi X, Fan Q, Zhang Y, Liang K, Wan L, Zhou Q, Li Y. 2020. Structural mechanism of phospholipids translocation by MlaFEDB complex. 12. Cell Res 30:1127–1135.

49. Powers MJ, Simpson BW, Trent MS. 2020. The Mla pathway in *Acinetobacter baumannii* has no demonstrable role in anterograde lipid transport. eLife 9:e56571.

50. Persky NS, Ferullo DJ, Cooper DL, Moore HR, Lovett ST. 2009. The ObgE/CgtA GTPase influences the stringent response to amino acid starvation in *Escherichia coli*. Mol Microbiol 73:253–266.

51. Manat G, Roure S, Auger R, Bouhss A, Barreteau H, Mengin-Lecreulx D, Touzé T. 2014. Deciphering the metabolism of undecaprenyl-phosphate: the bacterial cell-wall unit carrier at the membrane frontier. Microb Drug Resist 20:199–214.

52. Wijers CDM, Pham L, Menon S, Boyd KL, Noel HR, Skaar EP, Gaddy JA, Palmer LD, Noto MJ. 2021. Identification of two variants of *Acinetobacter baumannii* strain ATCC 17978 with distinct genotypes and phenotypes. Infect Immun 89:e0045421.

53. Lundstedt E, Kahne D, Ruiz N. 2021. Assembly and maintenance of lipids at the bacterial outer membrane. Chem Rev 121:5098–5123.

54. Emms DM, Kelly S. 2015. OrthoFinder: solving fundamental biases in whole genome comparisons dramatically improves orthogroup inference accuracy. Genome Biol 16:157.

55. Emms DM, Kelly S. 2019. OrthoFinder: phylogenetic orthology inference for comparative genomics. Genome Biol 20:238.

56. Sperandeo P, Lau FK, Carpentieri A, De Castro C, Molinaro A, Dehò G, Silhavy TJ, Polissi A. 2008. Functional analysis of the protein machinery required for transport of lipopolysaccharide to the outer membrane of *Escherichia coli*. J Bacteriol 190:4460–4469.

57. Lonergan ZR, Nairn BL, Wang J, Hsu Y-P, Hesse LE, Beavers WN, Chazin WJ, Trinidad JC, VanNieuwenhze MS, Giedroc DP, Skaar EP. 2019. An *Acinetobacter baumannii*, zinc-regulated peptidase maintains cell wall integrity during immune-mediated nutrient sequestration. Cell Rep 26:2009–2018.e6.

58. Dodbele S, Martinez CD, Troutman JM. 2014. Species differences in alternative substrate utilization by the antibacterial target undecaprenyl pyrophosphate synthase. Biochemistry 53:5042–5050.

59. Reid AJ, Scarbrough BA, Williams TC, Gates CE, Eade CR, Troutman JM. 2020. General utilization of fluorescent polyisoprenoids with sugar selective phosphoglycosyltransferases. Biochemistry 59:615–626.

60. Ragland SA, Criss AK. 2017. From bacterial killing to immune modulation: recent insights into the functions of lysozyme. PLoS Pathog 13:e1006512.

61. Malinverni JC, Silhavy TJ. 2009. An ABC transport system that maintains lipid asymmetry in the Gram-negative outer membrane. Proc Natl Acad Sci 106:8009–8014.

62. Jorgenson MA, Kannan S, Laubacher ME, Young KD. 2016. Dead-end intermediates in the enterobacterial common antigen pathway induce morphological defects in *Escherichia coli* by competing for undecaprenyl phosphate. Mol Microbiol 100:1–14.

63. Touzé T, Tran AX, Hankins JV, Mengin-Lecreulx D, Trent MS. 2008. Periplasmic phosphorylation of lipid A is linked to the synthesis of undecaprenyl phosphate. Mol Microbiol 67:264–277.

64. Loewen PC, Alsaadi Y, Fernando D, Kumar A. 2014. Genome sequence of an extremely drug-resistant clinical isolate of *Acinetobacter baumannii* strain AB030. Genome Announc 2:e01035–14.

65. Fujisaki S, Takahashi I, Hara H, Horiuchi K, Nishino T, Nishimura Y. 2005. Disruption of the structural gene for farnesyl diphosphate synthase in *Escherichia coli*. J Biochem (Tokyo) 137:395–400.

66. Takahashi H, Aihara Y, Ogawa Y, Murata Y, Nakajima K-I, Iida M, Shirai M, Fujisaki S. 2018. Suppression of phenotype of *Escherichia coli* mutant defective in farnesyl diphosphate synthase by overexpression of gene for octaprenyl diphosphate synthase. Biosci Biotechnol Biochem 82:1003–1010.

67. MacCain WJ, Kannan S, Jameel DZ, Troutman JM, Young KD. 2018. A defective undecaprenyl pyrophosphate synthase induces growth and morphological defects that are suppressed by mutations in the isoprenoid pathway of *Escherichia coli*. J Bacteriol 200:e00255–18.

68. Allen CM. 1985. Purification and characterization of undecaprenylpyrophosphate synthetase. Methods Enzymol 110:281–299.

69. Fujihashi M, Zhang YW, Higuchi Y, Li XY, Koyama T, Miki K. 2001. Crystal structure of cis-prenyl chain elongating enzyme, undecaprenyl diphosphate synthase. Proc Natl Acad Sci U S A 98:4337–4342.

70. Ko TP, Huang CH, Lai SJ, Chen Y. 2018. Structure of undecaprenyl pyrophosphate synthase from *Acinetobacter baumannii*. Acta Crystallogr Sect F Struct Biol Commun 74:765–769.

71. Touzé T, Mengin-Lecreulx D. 2008. Undecaprenyl phosphate synthesis. EcoSal Plus 3.

72. Guo R-T, Ko T-P, Chen AP-C, Kuo C-J, Wang AH-J, Liang P-H. 2005. Crystal structures of undecaprenyl pyrophosphate synthase in complex with magnesium, isopentenyl pyrophosphate, and farnesyl thiopyrophosphate: roles of the metal ion and conserved residues in catalysis. J Biol Chem 280:20762–20774.

73. Heuston S, Begley M, Gahan CGM, Hill C. 2012. Isoprenoid biosynthesis in bacterial pathogens. Microbiology 158:1389–1401.

74. Concha N, Huang J, Bai X, Benowitz A, Brady P, Grady LC, Kryn LH, Holmes D, Ingraham K, Jin Q, Pothier Kaushansky L, McCloskey L, Messer JA, O’Keefe H, Patel A, Satz AL, Sinnamon RH, Schneck J, Skinner SR, Summerfield J, Taylor A, Taylor JD, Evindar G, Stavenger RA. 2016. Discovery and characterization of a class of pyrazole inhibitors of bacterial undecaprenyl pyrophosphate synthase. J Med Chem 59:7299–7304.

75. Thorpe JH, Wall ID, Sinnamon RH, Taylor AN, Stavenger RA. 2020. Cocktailed fragment screening by X-ray crystallography of the antibacterial target undecaprenyl pyrophosphate synthase from *Acinetobacter baumannii*. Acta Crystallogr Sect F Struct Biol Commun 76:40–46.

76. Maneval WE. 1941. Staining bacteria and yeasts with acid dyes. Stain Technol 16:13–19.

77. Kang KN, Kazi MI, Biboy J, Gray J, Bovermann H, Ausman J, Boutte CC, Vollmer W, Boll JM. 2021. Septal class A penicillin-binding protein activity and LD-transpeptidases mediate selection of colistin-resistant lipooligosaccharide-deficient *Acinetobacter baumannii*. mBio 12:10.1128/mbio.02185-20.

78. Compton LA, Johnson WC. 1986. Analysis of protein circular dichroism spectra for secondary structure using a simple matrix multiplication. Anal Biochem 155:155–167.

79. Manavalan P, Johnson WC. 1987. Variable selection method improves the prediction of protein secondary structure from circular dichroism spectra. Anal Biochem 167:76–85.

80. Reid AJ, Eade CR, Jones KJ, Jorgenson MA, Troutman JM. 2021. Tracking Colanic Acid Repeat Unit Formation from Stepwise Biosynthesis Inactivation in Escherichia coli. Biochemistry 60:2221–2230.

81. Bai J, Raustad N, Denoncourt J, van Opijnen T, Geisinger E. 2023. Genome-wide phage susceptibility analysis in *Acinetobacter baumannii* reveals capsule modulation strategies that determine phage infectivity. PLOS Pathog 19:e1010928.

82. Geisinger E, Mortman NJ, Dai Y, Cokol M, Syal S, Farinha A, Fisher DG, Tang AY, Lazinski DW, Wood S, Anthony J, van Opijnen T, Isberg RR. 2020. Antibiotic susceptibility signatures identify potential antimicrobial targets in the *Acinetobacter baumannii* cell envelope. Nat Commun 11:4522.

83. Ondov BD, Treangen TJ, Melsted P, Mallonee AB, Bergman NH, Koren S, Phillippy AM. 2016. Mash: fast genome and metagenome distance estimation using MinHash. Genome Biol 17:132.

84. Letunic I, Bork P. 2007. Interactive Tree Of Life (iTOL): an online tool for phylogenetic tree display and annotation. Bioinforma Oxf Engl 23:127–128.

