## Supplementary Information for "Genetic synergy in *Acinetobacter baumannii* undecaprenyl biosynthesis and maintenance of lipid asymmetry impacts outer membrane and antimicrobial resistance"

**This PDF includes:**

Figures S1 to S3

Tables S1 to S3

References

**Figure S1. Dilution spotting of 17978UN WT, Δ*mlaF,* and Δ*mlaF* suppressor strains.**

Overnight cultures were serially diluted in PBS before plating on an LB agar plate with or without 0.01% SDS + 0.125 mM EDTA for overnight incubation. Image is representative of 4 biological replicates from 2 independent experiments.

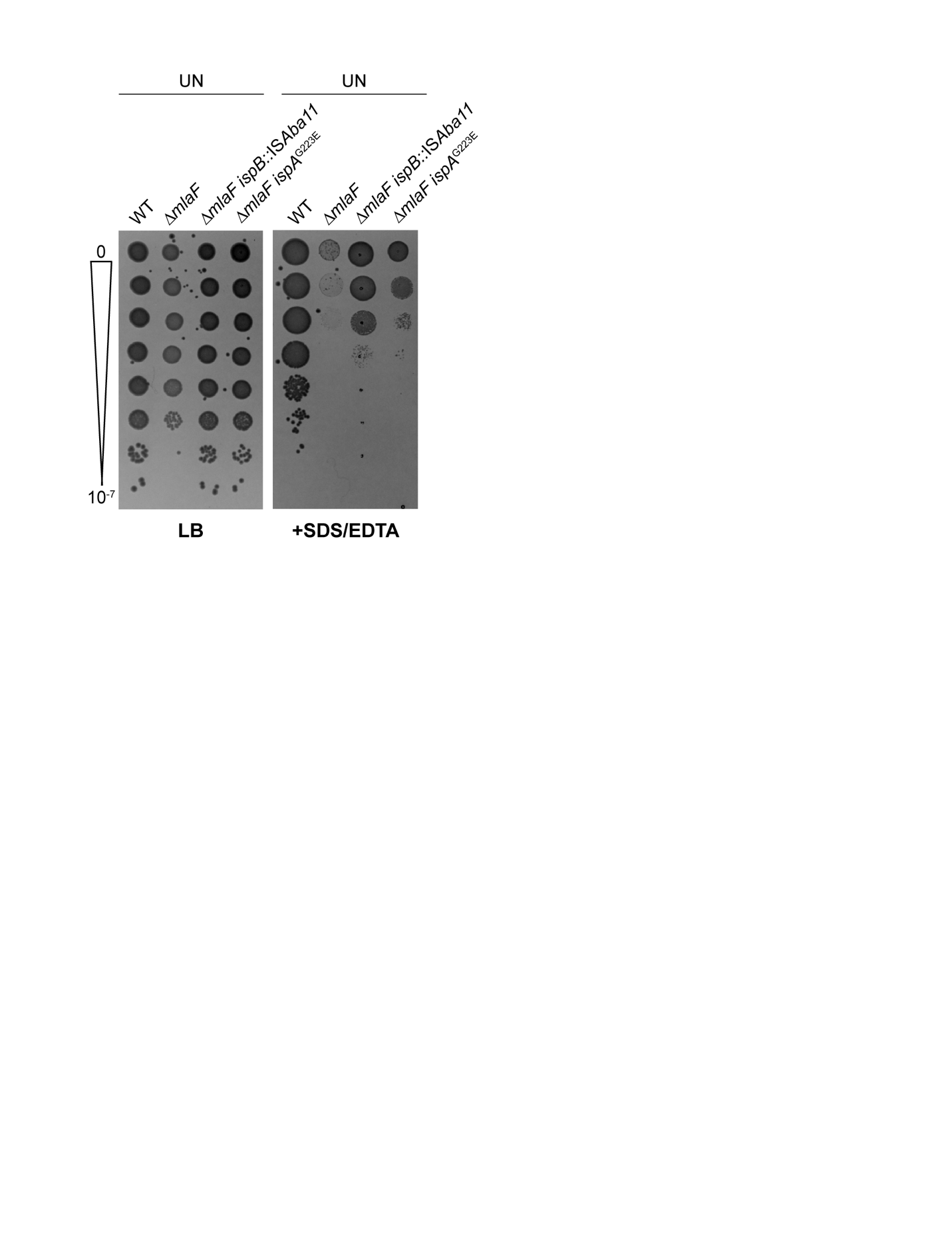

**Figure S2. Comparison of UppS^VU^ and UppS^UN^ protein secondary structure and 17978VU and 17978UN capsule, peptidoglycan, and LOS staining.**

**(A-B)** Purified UppS^VU^ and UppS^UN^ were subjected to circular dichroism analysis with a range of 190 nm to 210 nm.

**(C)** 17978VU and 17978UN WT and Δ*mlaF* strains were subjected to Maneval’s capsule stain. Scale bar = 5 μm. Images are representative of 4 biological replicates from 2 independent experiments.

**(D)** The peptidoglycan of 17978VU and 17978UN WT, Δ*mlaF, and* Δ*mlaF* with the opposite *uppS* allele was stained with NADA-green (3-[(7-Nitro-2,1,3-benzoxadiazol-4-yl)amino]-D-alanine hydrochloride). Scale bar = 5 μm. Images are representative of 2 biological replicates from 1 independent experiments.

**(E)** Raw image of silver stained LOS gel. Image is representative of 7 biological replicates from 3 independent experiments.

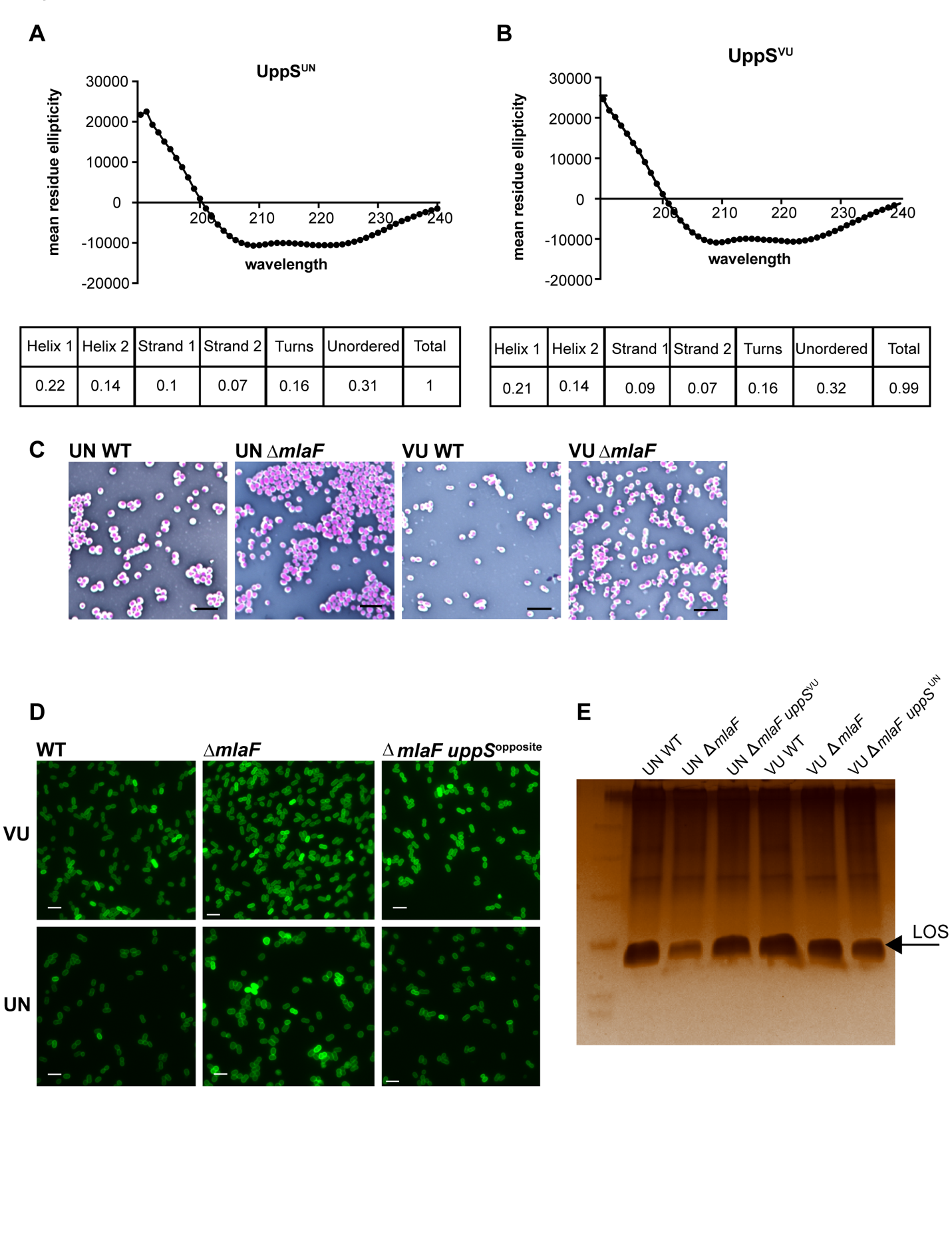

**Figure S3. Bacterial burdens from the murine lung infection model.**

**(A-D)** Bacterial CFU were enumerated 48 h after intranasal inoculation. n=5, medians are shown, and significance was determined by a Mann-Whitney test, * *p* < 0.05

**(E)** Growth curve in LB with and without lysozyme at 1 mg/mL. n=3, data are means +/- SEM.

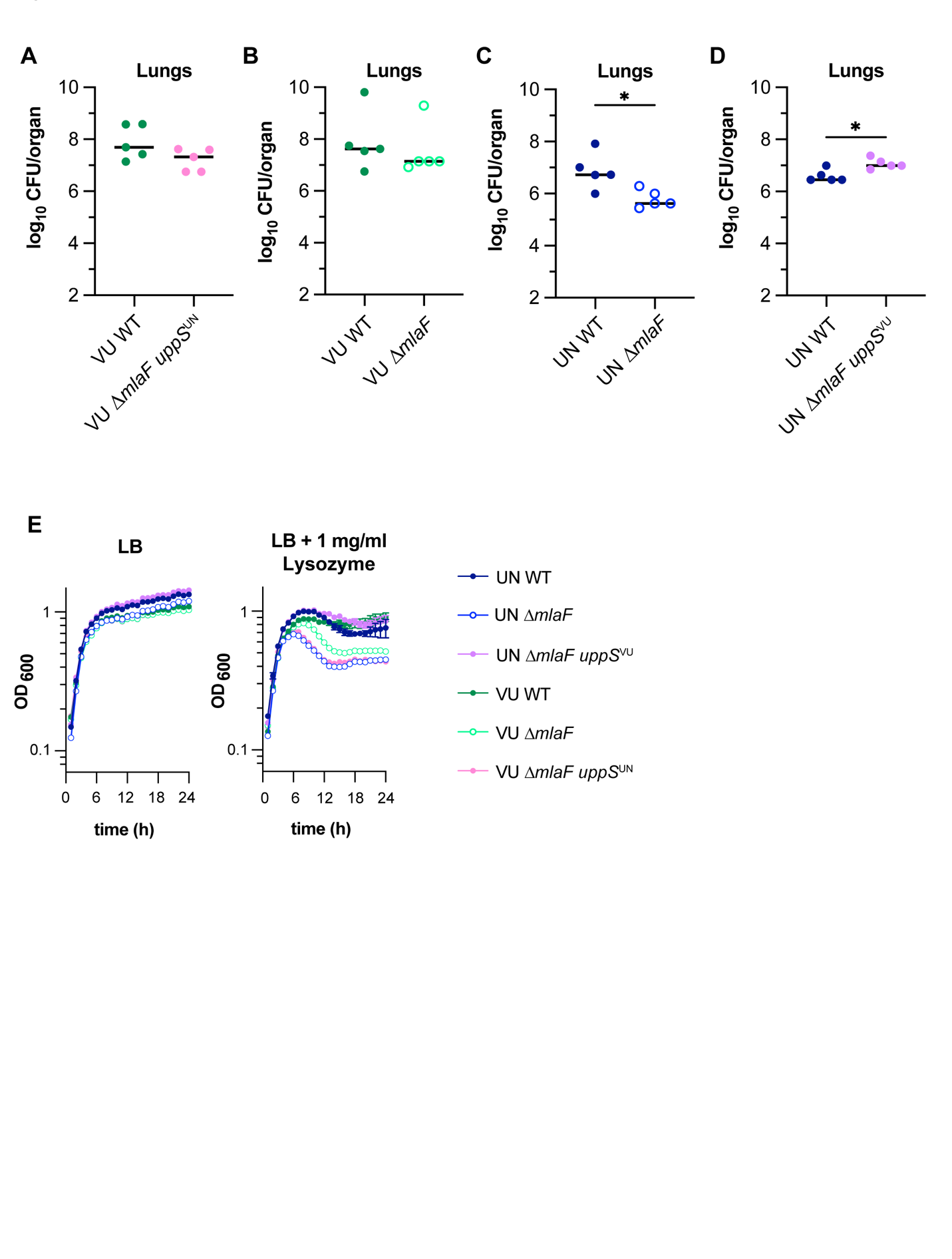

**Table S1. Strains**

| Strain | Source | Identifier |
| --- | --- | --- |
| *A. baumannii* 17978UN (WT) | ATCC | LP486 |
| *A. baumannii* 17978VU (WT) | ATCC | LP303 |
| *A. baumannii* 17978UN Δ*mlaF*(*A1S_3103*)::kan | Palmer et. al. 2020 (1) | LP39 |
| *A. baumannii* 17978VU Δ*mlaF*::kan | This manuscript | LP472 |
| *A. baumannii* 17978UN Δ*mlaF*::kan, *ACX60_03855*::IS*Aba11* | Palmer et al. 2020 (1) | LP338 |
| *A. baumannii* 17978VU Δ*mlaF*::kan, *ACX60_03855*::IS*Aba11* | This manuscript | LP485 |
| *A. baumannii* 17978VU, *obgE* I258 | This manuscript | LP477 |
| *A. baumannii* 17978VU Δ*mlaF*::kan, *obgE* I258 | This manuscript | LP484 |
| *A. baumannii* 17978VU, *lptD* V799 | This manuscript | LP521 |
| *A. baumannii* 17978VU Δ*mlaF*::kan, *lptD* V799 | This manuscript | LP522 |
| *A. baumannii* 17978UN, *uppS* M78 | This manuscript | LP628 |
| *A. baumannii* 17978VU, *uppS T*78 | This manuscript | LP523 |
| *A. baumannii* 17978UN Δ*mlaF*::kan, *uppS* M78 | This manuscript | LP629 |
| *A. baumannii* 17978VU Δ*mlaF*::kan, *uppS T*78 | This manuscript | LP540 |
| *A. baumannii* 17978UN Δ*mlaF*::kan, *ACX60_03855*::IS*Aba11*, *uppS* M78 | This manuscript | LP640 |
| *A. baumannii* 17978VU Δ*mlaF*::kan, *ACX60_03855*::IS*Aba11*, *uppS T*78 | This manuscript | LP622 |
| *A. baumannii* 17978UN Δ*mlaF*::kan, *uppS T*78, *ispA* (*ACX60_03710) G223E* | This manuscript | LP798 |

**Table S2. Plasmids**

| Use or Phenotype | Source | Name |
| --- | --- | --- |
| KnR; mobilization helper plasmid | Figurski and Helinski, 1979 (2) | pRK2013 |
| CarbR; allelic exchange vector with sucrose resistance | Hoang et al., 1998 (3) | pFLP2 |
| Contains an FRT-flanked kanamycin resistance gene | Datsenko and Wanner, 2000 (4) | pKD4 |
| CarbR; allelic exchange vector to delete *mlaF*; pFLP2-Δ*mlaF*::Kn | Palmer et. al. 2020 (1) | pLDP8 |
| CarbR; allelic exchange vector to swap *obgE* I258 allele to *obgE* N258; pFLP2-*obgE** | This manuscript | pLDP84 |
| CarbR; allelic exchange vector to swap *obgE* N258 allele to *obgE* I258; pFLP2-*obgE* | This manuscript | pLDP88 |
| CarbR; allelic exchange vector to swap *lptD* allele to *lptD* V799; pFLP2-*lptD* V799 | This manuscript | pLDP104 |
| CarbR; allelic exchange vector to swap *lptD* allele to *lptD F*799; pFLP2-*lptD* F799 | This manuscript | pLDP105 |
| CarbR; allelic exchange vector to swap *uppS* allele to *uppS* M78; pFLP2-*uppS*^M78^ | This manuscript | pLDP106 |
| CarbR; allelic exchange vector to swap *uppS* allele to *uppS*^T78^; pFLP2-*uppS*^T78^ | This manuscript | pLDP107 |
| CarbR; allelic exchange vector to regenerate the ispA* suppressor mutation (*ACX60_03710 G223E)*; pFLP2-ispA* | This manuscript | pLDP187 |
| CarbR; UppS^VU^ expression vector for protein purification | This manuscript | pLDP192 |
| CarbR; UppS^UN^ expression vector for protein purification | This manuscript | pLDP193 |

**Table S3. Oligonucleotides**

| Name | Sequence | Source | Name |
| --- | --- | --- | --- |
| pFLP2 fwd | tgaacggcaggtatatgtgatggg | IDT (Coralville, IA) | LP52 |
| pFLP2 rev | ccatgattacgaattcgagc | IDT (Coralville, IA) | LP54 |
| pKNOCKseq_fwd | gcgcttttgaagctaattcg | IDT (Coralville, IA) | LP354 |
| pKNOCKseq rev | atttcacttatctggttggcc | IDT (Coralville, IA) | LP355 |
| pFLP2_ObgE_fwd | aaaggatcgatcctctagagTGTTCATCCTTAAAGCTAATTTG | IDT (Coralville, IA) | HN1 |
| pFLP2_ObgE_rev | tacgaattcgagctcggtacCTGTTAAAATAATTAATAAATCAGCATGTAC | IDT (Coralville, IA) | HN2 |
| Up_ObgE_fwd | Ctctgttcagcacactaaagc | IDT (Coralville, IA) | HN3 |
| Dn_ObgE_fwd | Caccaccaccagccatttc | IDT (Coralville, IA) | HN4 |
| ObgE*_seq_fwd | gtacccgttggtacaacaattg | IDT (Coralville, IA) | HN5 |
| ObgE*_seq_rev | gattcgtttacactcactgagc | IDT (Coralville, IA) | HN6 |
| ObgE_MutSite_fwd | gttcagatccggcccataa | IDT (Coralville, IA) | HN7 |
| ObgE*_MutSite_fwd | gttcagatccggcccatat | IDT (Coralville, IA) | HN8 |
| pFLP2_LptD_FWD | GGTTAAAAAGGATCGATCCTCTAGAGGATCattctgcacaagaaagtagcggtacgcgta | IDT (Coralville, IA) | HN23 |
| pFLP2_LptD_rev | TGACCATGATTACGAATTCGAGCTCGGTACaatcttcacccgcttttaatcgcttgtaaa | IDT (Coralville, IA) | HN24 |
| pFLP2_UppS_fwd | GATCGATCCTCTAGAGGATCtttaacggcaatttgcttatg | IDT (Coralville, IA) | HN25 |
| pFLP2_UppS_rev | TACGAATTCGAGCTCGGTACttaccgaatttacggcctac | IDT (Coralville, IA) | HN26 |
| Up_LptD_fwd | gatgcagacaagccatatg | IDT (Coralville, IA) | HN27 |
| Dn_LptD_rev | cctacaggaatttcctgc | IDT (Coralville, IA) | HN28 |
| Up_UppS_fwd | gaaagcgaccaaagtagac | IDT (Coralville, IA) | HN29 |
| Dn_UppS_rev | caagaatagaagttgttgaaccc | IDT (Coralville, IA) | HN30 |
| UppS_T78mutsite_fwd | caatatgaagtcgatctgcttac | IDT (Coralville, IA) | HN31 |
| UppS_M78mutsite_fwd | caatatgaagtcgatctgcttat | IDT (Coralville, IA) | HN32 |
| UppS_seq_fwd | ctattactgctgattcaatccg | IDT (Coralville, IA) | HN33 |
| LptD_F799mutsite_fwd | cgtctttacttgaaaatcgct | IDT (Coralville, IA) | HN34 |
| LptD_V799mutsite_fwd | gtctttacttgaaaatcgcg | IDT (Coralville, IA) | HN35 |
| LptD_seq_fwd | gctattattttgaagatcgcc | IDT (Coralville, IA) | HN36 |
| pFLP2-ispA*_f | AAAAGGATCGATCCTCTAGAGGATCggctggtggtggagaacg | IDT (Coralville, IA) | HN85 |
| pFLP2-ispA*_r | ATGACCATGATTACGAATTCGAGCTgcaattttagaagcagatggtcgtt | IDT (Coralville, IA) | HN86 |
| dn_dn_ispA_r | caacgtcgtgctacagttac | IDT (Coralville, IA) | HN87 |
| up_up_ispA*_f | cagcagcattagcacgttttg | IDT (Coralville, IA) | HN88 |
| ispA*seq_f | cttgtatggacaatgatttgcttc | IDT (Coralville, IA) | HN89 |
| pET15b-UppS-fwd | ttcgggctttgttagcagccggatcttataatttctcgattttctcttgc | IDT (Coralville, IA) | HN94 |
| pET15b-UppS-rev | ggcctggtgccgcgcggcagccatatgaccgattcagaagagtat | IDT (Coralville, IA) | HN95 |

References

1.	Palmer LD, Minor KE, Mettlach JA, Rivera ES, Boyd KL, Caprioli RM, Spraggins JM, Dalebroux ZD, Skaar EP. 2020. Modulating isoprenoid biosynthesis increases lipooligosaccharides and restores *Acinetobacter baumannii* resistance to host and antibiotic stress. Cell Rep 32:108129.

2.	Figurski DH, Helinski DR. 1979. Replication of an origin-containing derivative of plasmid RK2 dependent on a plasmid function provided in trans. Proc Natl Acad Sci U S A 76:1648–1652.

3.	Hoang TT, Karkhoff-Schweizer RR, Kutchma AJ, Schweizer HP. 1998. A broad-host-range Flp-FRT recombination system for site-specific excision of chromosomally-located DNA sequences: application for isolation of unmarked *Pseudomonas aeruginosa* mutants. Gene 212:77–86.

4.	Datsenko KA, Wanner BL. 2000. One-step inactivation of chromosomal genes in *Escherichia coli* K-12 using PCR products. Proc Natl Acad Sci U S A 97:6640–6645.
